# Immunological Profiling in Knee Osteoarthritis: Treg Dysfunction as Key Driver of Pain

**DOI:** 10.1101/2024.10.12.618016

**Authors:** Marie Binvignat, Johanna Dubois, Maria Marco Salvador, Paul Stys, Fabien Pitoiset, Alexandra Roux, Michèle Barbié, Signe Hassler, Roberta Lorenzon, Claire Ribet, Vanessa Mhanna, Hélène Vantomme, Leslie Adda, Pierre Barennes, Nicolas Coatnoan, Kenz Le Gouge, Caroline Aheng, Alice Courties, Lise Minssen, Atul J. Butte, Adrien Six, Michelle Rosenzwajg, Nicolas Tchitchek, Francis Berenbaum, David Klatzmann, Encarnita Mariotti-Ferrandiz, Jérémie Sellam

## Abstract

Pain is the hallmark symptom of osteoarthritis (OA) and its biological drivers remain poorly understood. While the role of innate immunity in OA has been extensively studied, the involvement of adaptive immunity, in particular regulatory T cells (Tregs), is not well understood. Using a comprehensive multi-omic approach on the peripheral blood from 46 knee OA patients with similar radiographic stage, including deep immunophenotyping, cytokine profiling, transcriptomic and T-cell receptor analysis on sorted CD4 Tregs and effector T cells (Teff), we identified an immunological signature associated with OA-related pain. Cytokines promoting Treg expansion and activation (with increases of sIL2-RA, sTNFR1, sTNFR2) were correlated with the Western Ontario and McMaster Universities Arthritis Index (WOMAC) pain subscore, suggesting a potential Treg dysfunction. Nineteen T cell subsets were correlated with WOMAC pain. Notably, we found a negative correlation of cell subsets associated with Treg expansion and activation (FoxP3+CTLA4+, CD4+CD57+, Treg CD95+, CD4 Treg CD45RA-). Differential gene expression analysis between patients with low and high WOMAC pain intensity (threshold ≥ 40/100) revealed an upregulation of inflammasome-related genes such as *IL1RL1, IL31RA, IFITM3, NLRP3, IFNG* in Tregs. Functional enrichment analysis highlighted an overrepresentation of innate immune response, IL-8, and interferon activation pathways suggesting a pro-inflammatory state in Tregs of patients with high pain intensity. Collectively, our systems immunology approach highlights multiple associations between Treg dysfunctionality and OA-related pain, providing new insights into the adaptive immune system’s contribution to OA-related pain.

## INTRODUCTION

Osteoarthritis (OA) is the most prevalent joint disorder, affecting approximately 8% of the global population (*1*, *2*). It is a leading cause of disability, comorbidities, and societal burden and knee OA is accounting for nearly 85% of the disease impact (*3–6*). Pain is the primary symptom of OA and its alleviation represents the main goal for both patients and physicians, serving as the principal driver of therapeutic decisions (*7*, *8*). Despite its high prevalence, the pathophysiology underlying OA-related pain remains poorly understood, and available therapeutic options are limited and often associated with safety concerns, particularly in the context of the ongoing opioid crisis (*9*, *10*). In recognition of this significant unmet medical need, the Food and Drug Administration (FDA) has classified OA as a serious disease, thereby facilitating the development and expedited review of novel therapeutic agents (*11*, *12*). Notably, novel pharmacological therapeutic targets are needed, since up to 20% of patients still experience pain after total knee replacement (*13*). Magnetic resonance imaging (MRI) studies indicated that nociceptive pain was driven by synovitis or joint effusion and bone marrow lesions (BML) in patients with knee OA-related pain although the immune and inflammatory mechanisms behind remain not completely understood (*14–18*). OA-related pain, while seemingly straightforward, is complex and heterogeneous with not only nociceptive but also neuropathic-like, and nociplastic components (*19*, *20*). As such, a deeper understanding of these pain mechanisms and the identification of distinct pain phenotypes and endotypes are essential for advancing effective therapeutic strategies (*20–22*).

Among all biological mechanisms involved in OA, local and systemic low-grade inflammation has emerged as a key driver of pain, with the innate immune response playing a central role in mediating OA-related nociception (*23*). Proinflammatory cytokines are implicated in pain through mechanisms such as cartilage degradation, synovial inflammation, bone remodeling and direct activation of nociceptors (*24*, *25*). Additionally, recent studies in murine models have shown that M1-like macrophages accumulate in the dorsal root ganglia (DRG), further highlighting the contribution of innate immunity to OA-related pain (*26*). While extensive research has focused on innate immune mechanisms, the role of adaptive immunity in OA remains underexplored. Emerging evidence suggests a potential involvement of adaptive immune cells, including T cells and especially regulatory T cells (Tregs), in OA-related pain and inflammation. Tregs, which are key players of peripheral tolerance and immune regulation (*27*), could influence OA pathophysiology, with reduced Treg levels being linked to increased knee OA-related pain (*28*). In addition, Interleukin-2 (IL-2), essential for Treg development and function, has been suggested as a potential biomarker in OA (*29*, *30*). These preliminary data suggest that adaptive immunity may represent a novel therapeutic target in OA, warranting further investigation into its role in disease progression and pain management (*31*).

In this study, we aimed to provide a comprehensive and systematic immunological profiling of OA-related pain through extensive analysis of peripheral blood. Our primary objective was to identify an immunological signature associated with OA-related pain using a multi-omic approach, including deep immunophenotyping, cytokine profiling, as well as transcriptomic and T-cell receptor (TCR) sequencing analysis on sorted Tregs and effector T cells. Additionally, our secondary objective was to explore potential associations between immunological markers linked to OA-related pain and MRI markers of BML and knee effusion - synovitis.

## RESULTS

### Transimmunom cohort characteristics and study design

We enrolled 46 patients from the Transimmunom study (*32*) with symptomatic knee OA. Knee OA was defined according to the ACR criteria (*33*), and patients had to have a radiographic severity with a Kellgren-Lawrence score of 2 or 3 to avoid doubtful (Kellgren-Lawrence score of 1) or end-stage OA (Kellgren score 4) in which pain can be biomechanically induced (*34*) (**Table S1**). This study included 65.2% of women (N=30) with a mean age of 64.8 years (SD=9.9) and a mean BMI of 29.1 (SD=7.1) (**Table 1**). Western Ontario and McMaster Universities Osteoarthritis Index (WOMAC) pain subscore was used to assess OA-related pain with a mean score of 44.2/100 (12.7). Neuropathic pain was defined using the *Douleur Neuropathique 4* (DN4) score (used to assess neuropathic-like pain) with a mean value of 2.1/10 (1.93), and depression was evaluated using the Hospital Anxiety and Depression Scale (HADS) with a mean score of 13.3/42 (7.5). Forty-one patients underwent a standardized knee MRI with semi-quantification of knee effusion-synovitis and BML according to the Knee Osteoarthritis Scoring System (KOSS) on the most symptomatic knee (*35*). Among these patients, twenty-five exhibited BML (≥ 5 mm), and sixteen displayed moderate to severe knee effusion-synovitis. Patients were also stratified in low (N=16) and high pain intensity groups (N=30) using a WOMAC pain subscore threshold of 40/100 corresponding to the patient acceptable symptom state (PASS) (*36*). There was no significant difference in age, BMI, sex distribution, Kellgren-Lawrence score of the target knee, or OA duration between patients with low and high pain intensity (**Table 1**, **Fig. S1)**. For each patient, we performed deep immunophenotyping on peripheral blood using 12 different flow cytometry panels following 269 cell populations. We also analyzed 62 serum cytokines, chemokines and soluble receptors and conducted TCR and mRNA sequencing on sorted CD4+CD25+CD127low Treg and CD4+CD25+CD127+ Teff cells. The study design is summarized in **Fig 1A**.

**Table 1.**
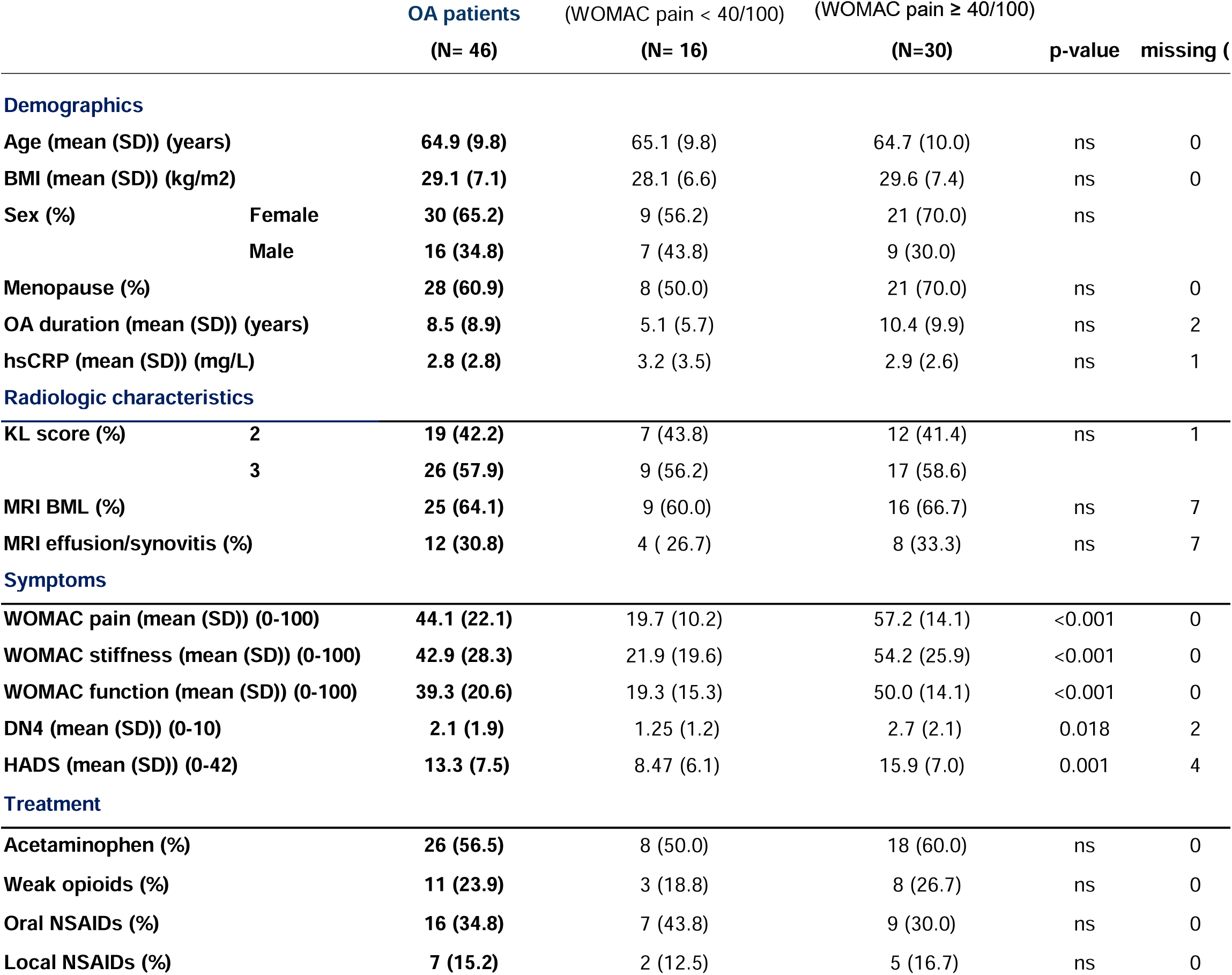
Clinical characteristic of OA patients with low and high WOMAC pain score (≥ 40/100). Continuous variables were expressed as mean and standard deviation (SD) in each subgroup, while categorical variables were expressed as percentages. A Student’s t-test and chi-square test were applied to assess the differences between patients with low and high levels of pain using a WOMAC pain score cut-off of 40/100. A significant p-value was defined with a threshold of ≤ 0.05. Missing values are expressed as absolute number (N).

| | | **OA patients**<br>**(N= 46)** | (WOMAC pain < 40/100)<br>**(N= 16)** | (WOMAC pain $\geq 40/100$)<br>**(N=30)** | **p-value** | **missing (N)** |
| --- | --- | --- | --- | --- | --- | --- |
| **Demographics** |  |  |  |  |  |  |
| **Age (mean (SD)) (years)** |  | **64.9 (9.8)** | 65.1 (9.8) | 64.7 (10.0) | ns | 0 |
| **BMI (mean (SD)) (kg/m<sup>2</sup>)** |  | **29.1 (7.1)** | 28.1 (6.6) | 29.6 (7.4) | ns | 0 |
| **Sex (%)** | **Female** | **30 (65.2)** | 9 (56.2) | 21 (70.0) | ns |  |
|  | **Male** | **16 (34.8)** | 7 (43.8) | 9 (30.0) |  |  |
| **Menopause (%)** |  | **28 (60.9)** | 8 (50.0) | 21 (70.0) | ns | 0 |
| **OA duration (mean (SD)) (years)** |  | **8.5 (8.9)** | 5.1 (5.7) | 10.4 (9.9) | ns | 2 |
| **hsCRP (mean (SD)) (mg/L)** |  | **2.8 (2.8)** | 3.2 (3.5) | 2.9 (2.6) | ns | 1 |
| **Radiologic characteristics** |  |  |  |  |  |  |
| **KL score (%)** | **2** | **19 (42.2)** | 7 (43.8) | 12 (41.4) | ns | 1 |
|  | **3** | **26 (57.9)** | 9 (56.2) | 17 (58.6) |  |  |
| **MRI BML (%)** |  | **25 (64.1)** | 9 (60.0) | 16 (66.7) | ns | 7 |
| **MRI effusion/synovitis (%)** |  | **12 (30.8)** | 4 (26.7) | 8 (33.3) | ns | 7 |
| **Symptoms** |  |  |  |  |  |  |
| **WOMAC pain (mean (SD)) (0-100)** |  | **44.1 (22.1)** | 19.7 (10.2) | 57.2 (14.1) | <0.001 | 0 |
| **WOMAC stiffness (mean (SD)) (0-100)** |  | **42.9 (28.3)** | 21.9 (19.6) | 54.2 (25.9) | <0.001 | 0 |
| **WOMAC function (mean (SD)) (0-100)** |  | **39.3 (20.6)** | 19.3 (15.3) | 50.0 (14.1) | <0.001 | 0 |
| **DN4 (mean (SD)) (0-10)** |  | **2.1 (1.9)** | 1.25 (1.2) | 2.7 (2.1) | 0.018 | 2 |
| **HADS (mean (SD)) (0-42)** |  | **13.3 (7.5)** | 8.47 (6.1) | 15.9 (7.0) | 0.001 | 4 |
| **Treatment** |  |  |  |  |  |  |
| **Acetaminophen (%)** |  | **26 (56.5)** | 8 (50.0) | 18 (60.0) | ns | 0 |
| **Weak opioids (%)** |  | **11 (23.9)** | 3 (18.8) | 8 (26.7) | ns | 0 |
| **Oral NSAIDs (%)** |  | **16 (34.8)** | 7 (43.8) | 9 (30.0) | ns | 0 |
| **Local NSAIDs (%)** |  | **7 (15.2)** | 2 (12.5) | 5 (16.7) | ns | 0 |

**Fig. 1.**
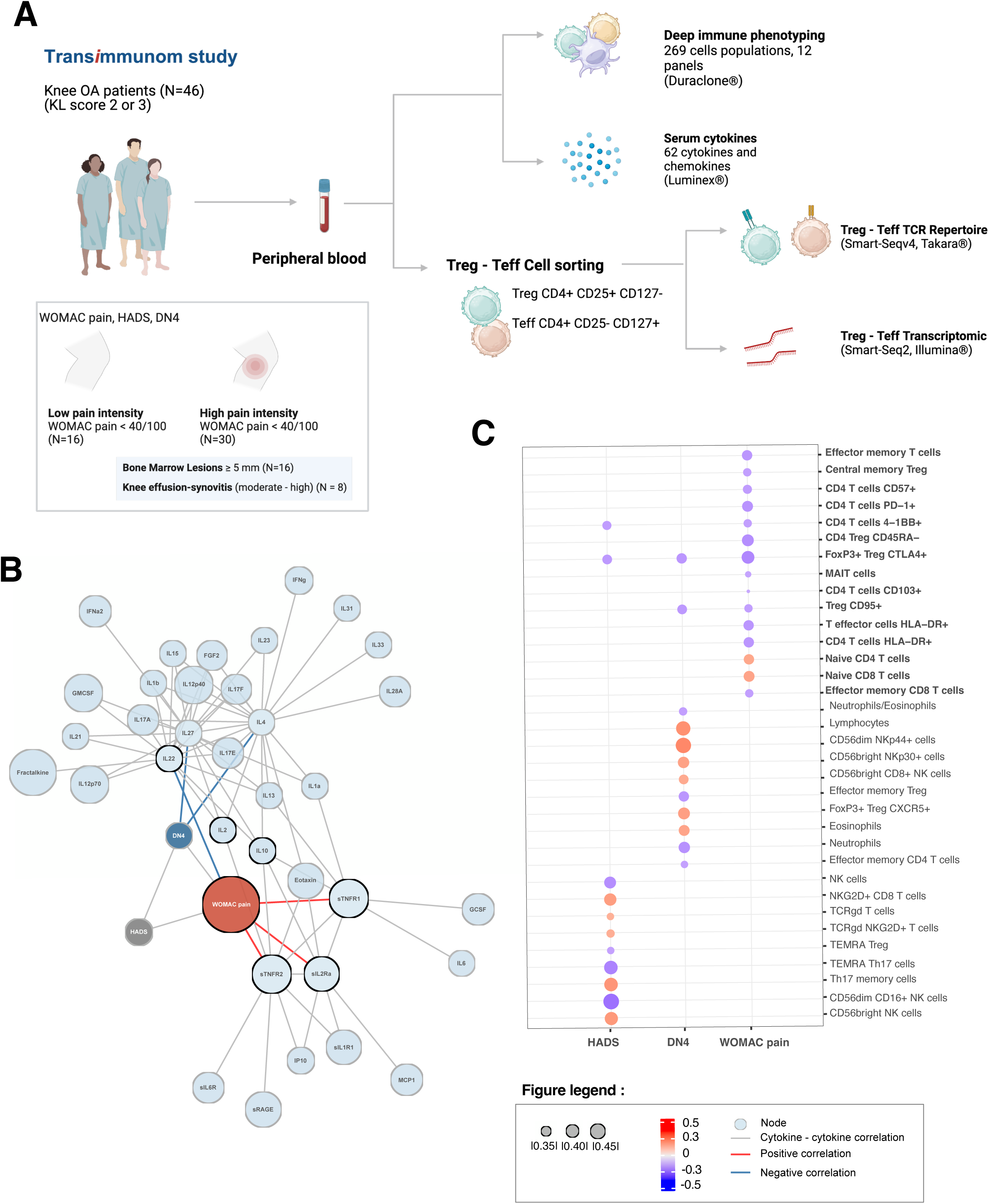
Deep Immunophenotyping and cytokines analysis identify a specific T cell signature of OA-related pain. A. Transimmunom study design. B. Correlation network between symptoms (WOMAC pain, DN4 and HADS) and cytokines, negative correlation are in blue and positive correlation in red, cytokines-cytokines correlation and symptoms-symptoms correlation are in gray |r| ≥ 0.2 p≤ 0.05. C. Correlation dot-plot with significant subset associated with HADS, DN4 and WOMAC pain using a Spearman correlation (N=42) and positive correlation are in red, negative correlation in blue |r| ≥0.2, p<0.05

### sIL2-RA, sTNFR1, sTNFR2, IL-22 were associated with WOMAC pain

In order to investigate the modulation of the immune response in relation with pain in OA, we first analyzed 62 cytokine, chemokine and soluble receptor levels from serum samples (**Table S2**) and their correlation with WOMAC pain, DN4 and HADS using non parametric Spearman correlation and we applied an absolute correlation of threshold |r| ≥ 0.2, a p-value threshold ≤ 0.05 (N= 44 patients). We found a positive correlation between sIL2RA and WOMAC pain (r=0.31, p=0.042). Similarly, sTNFR1 and sTNFR2 were positively correlated with WOMAC pain (r=0.33, p=0.028 and r=0.31, p=0.031, respectively) while IL-22 was negatively correlated with WOMAC pain (r=-0.33, p=0.028) (**Fig. 1B, Fig. S1, Table S3**). DN4 scores were negatively correlated with IL-27 and IL-4 levels (r=-0.35, p=0.021 and r=-0.33, p=0.028, respectively), and no significant correlations were found between HADS scores and any cytokine, chemokine and soluble receptor levels. Altogether, these results highlight several cytokines and soluble receptors that may be involved in immune regulation and inflammation specifically associated with pain intensity (ie WOMAC pain) in knee OA patients and suggest a potential role of Tregs through higher IL2-RA levels.

### WOMAC pain is associated with Treg expansion and activation

We then analyzed the modulation of the immune response at the level of the immune cell subsets by performed a deep immunophenotyping using 12 standardized panels of flow cytometry (*37*, *38*) on 46 knee OA patients. With this approach, 269 cell parameters were systematically and simultaneously measured on each blood sample (**Fig.S2, Table S4**). We first calculated the Spearman’s correlations of the cell subset percentage with WOMAC pain score, DN4, and HADS **(Fig. 1C, Fig S3-S4, Table S5**). Nineteen cell populations are associated with WOMAC pain, 12 with DN4, and 11 with HADS (|r| > 0.2, p ≤ 0.05). Interestingly, the cell populations correlated with OA-related pain intensity were distinct from those associated with neuropathic-like pain and depression, suggesting that specific immune cell populations may be uniquely linked to OA-related pain. In addition, WOMAC pain was predominantly associated with CD4 T cells and Tregs while DN4 and HADS were more likely associated with CD8+ T cells and NK cells. In particular, WOMAC pain was negatively correlated with several cell subsets associated with Treg and T cell activation including Treg CD95+ (r = −0.33, p =0.028), Treg FoxP3+ CTLA4+ (r = −0.41, p = 0.006), CD4 Treg CD45RA-(r = −0.39, p = 0.009) and CD4+ CD57+ T cells (r = −0.34, p = 0.023). Altogether, our results identified a distinct set of cell subsets correlated with pain intensity in OA patients, predominantly mediated by the CD4+ Treg cell subsets.

### Upregulation of inflammasome-related genes in Tregs linked with high OA-related pain intensity

To further investigate the role of Treg cells in OA-related pain, we compared mRNA transcriptomics on sorted Tregs and Teff cells between patients with low and high pain intensity (WOMAC pain ≥ 40/100). CD4+ T cells were extracted from peripheral blood and subsequently sorted into Treg (CD4+CD25+CD127-/low) and Teff (CD4+CD25-CD127+) subsets. We performed mRNA transcriptomic using next generation sequencing (NGS) on those sorted cells. Differential gene expression and functional analysis were conducted between patients with low and high pain intensity. We identified 69 differentially expressed genes in Tregs between patients with low pain intensity (N=6) and patients with high pain intensity (N=28) (|log2 FC | ≥ log2(1.6), p ≤ 0.01, baseMean expression ≥ 8) (Fig.2, Fig S5-7, Table S6-7). Among these, 46 genes were upregulated, and 23 genes were downregulated. We observed an upregulation of proinflammatory genes and genes related to the inflammasome and interferon pathways with OA-related pain including *IL1RL1*, *IL31RA*, *IFITM3*, *IFI44L*, and *NLRP3*. We also identified 60 differentially expressed genes in Teff between patients with low pain intensity (N=14) and those with high pain intensity (N=28), comprising 11 upregulated and 49 downregulated genes (|log2 FC | ≥ log2(1.6), p ≤ 0.01, baseMean expression ≥ 0.8). Genes associated with bone remodeling, such as TNFRS11A (RANK) and SECTM1, as well as genes linked to NK and CD8 cytotoxicity, including CDKN1C and KLRC4-KLRK1, were downregulated in patients with high pain intensity. Additionally, pro-inflammatory genes such as TNFSF13, TNFSF13B, KYNU, IFI30, and NFIL3 showed downregulation.

**Fig. 2.**
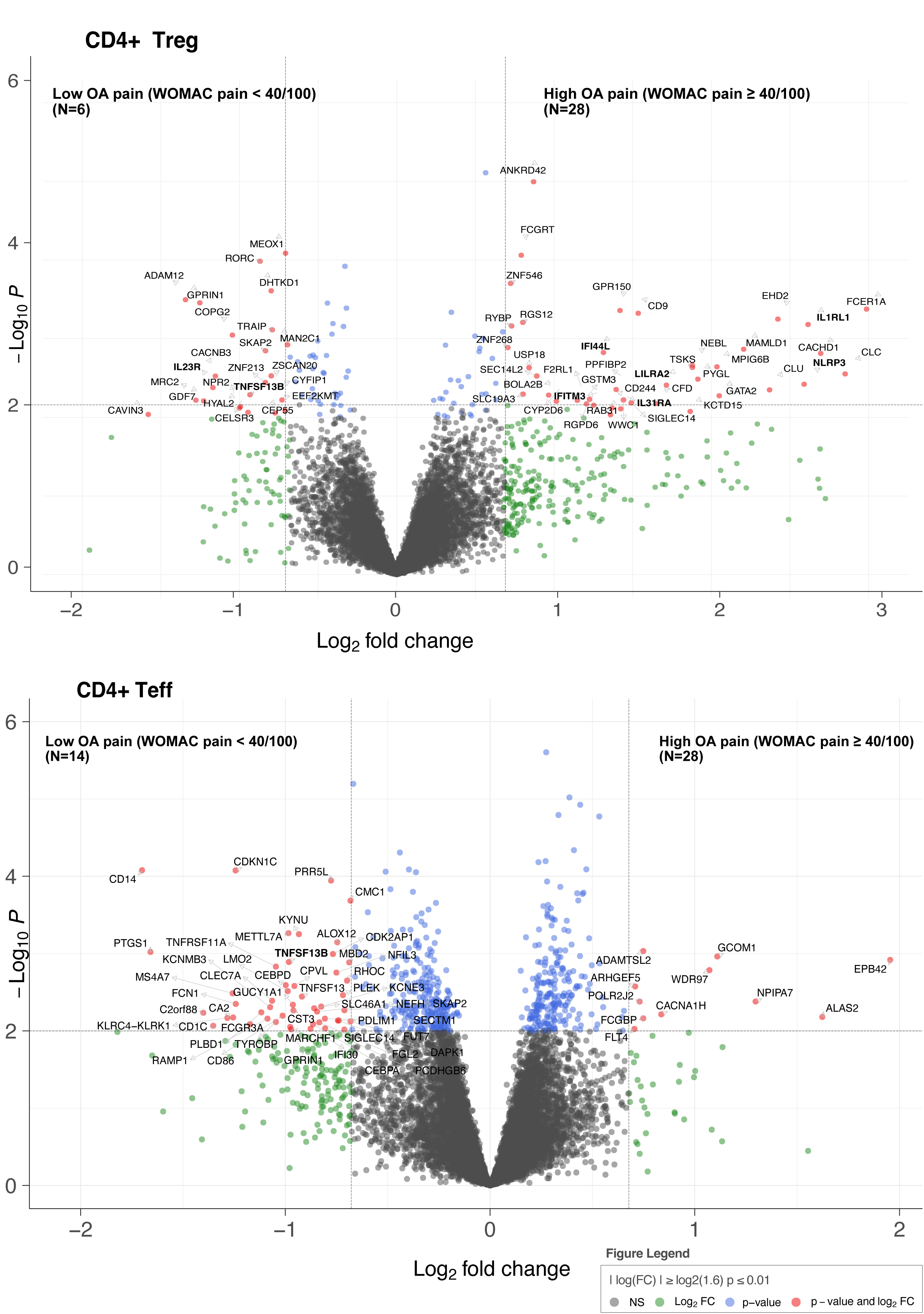
Treg and Teff transcriptomic analysis between patients with low and high pain intensity. CD4+ Tregs **and Teff** Volcano plots of differentially expressed genes between patients with low (N=6, N =14) and high pain intensity (N=28) (|log2 FC | ≥ log2(1.6), p-value ≤ 0.01, baseMean expression ≥ 8).

Over-representation and GO pathways analysis in Tregs showed an enrichment of pro-inflammatory pathways related to innate immune activation, IL-8 and IFN-gamma cytokine production, and in Teff pathways associated with immune cellular response, as well as leukocyte-mediated cytotoxicity, regulation of cell killing, and natural killer-mediated cytotoxicity (p<0.05, |log2 FC | ≥ log2(1.6), baseMean expression ≥ 0.8, FDR ≤ 0.05) (Fig. 3). Gene-concept network analysis showed an important contribution of NLRP3 and IL1RL1 in the positive regulation of cytokine production and IL1RL1 in myeloid leukocyte activation in Tregs while KLRC4-KLRK1 was highlighted for its significant role in leukocyte-mediated cytotoxicity, innate immune response activation, signal transduction and surface receptor signaling pathways in Teffs (**Fig. 4**). Altogether, these results highlight a dysregulation of peripheral blood Tregs toward a proinflammatory phenotype in patients with high levels of pain. In contrast, Teff displayed reduced effector functions in patients with high pain intensity.

**Fig. 3.**
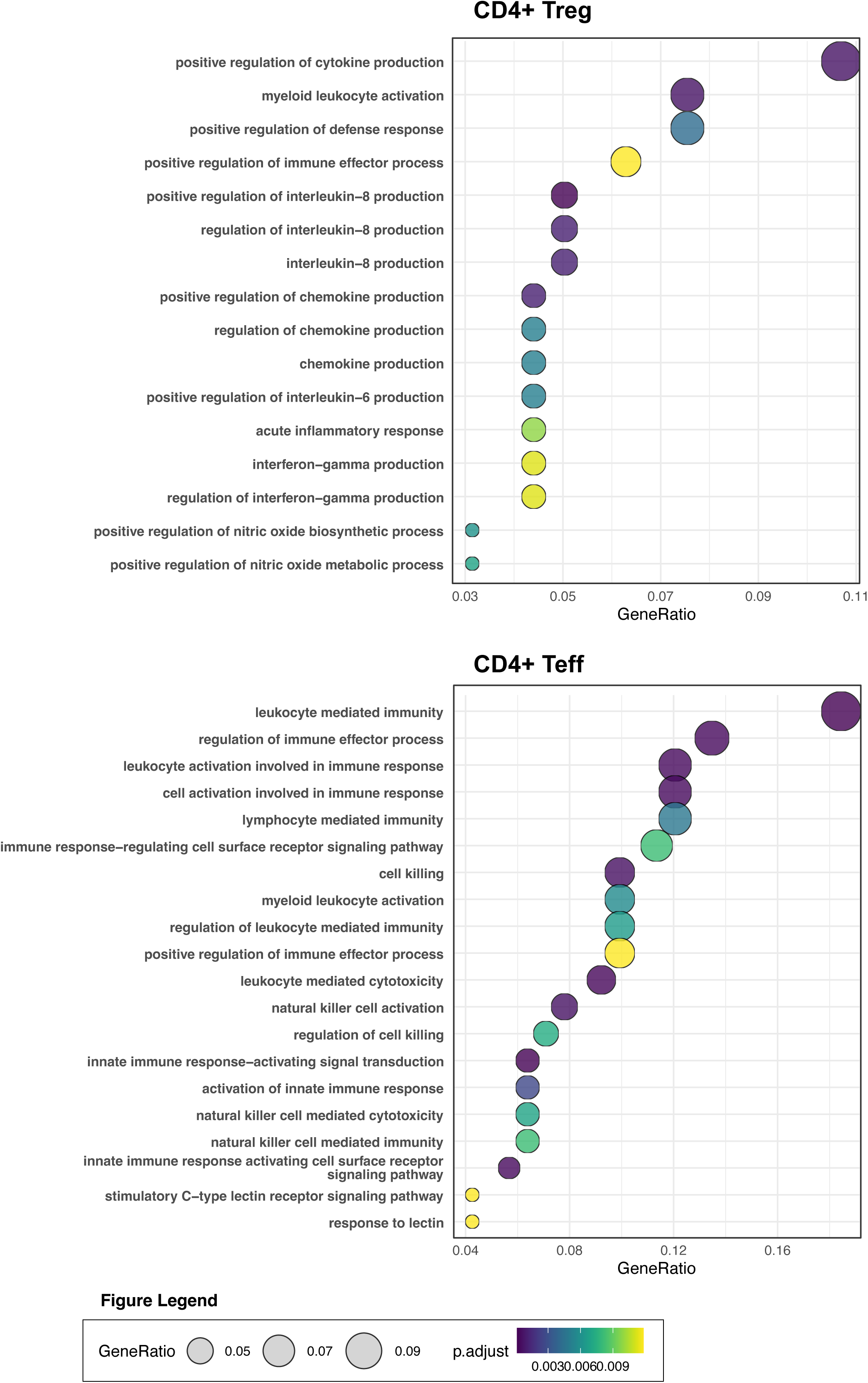
CD4+ Tregs and Teff functional analysis. CD4+ Treg and Teff dot plots of over-representation analysis and enriched GO pathway ordered by gene ratio (N=42) (|log2 FC | ≥ log2(1.6), p-value ≤ 0.05, FDR ≤0.01, baseMean expression ≥ 8

**Fig. 4.**
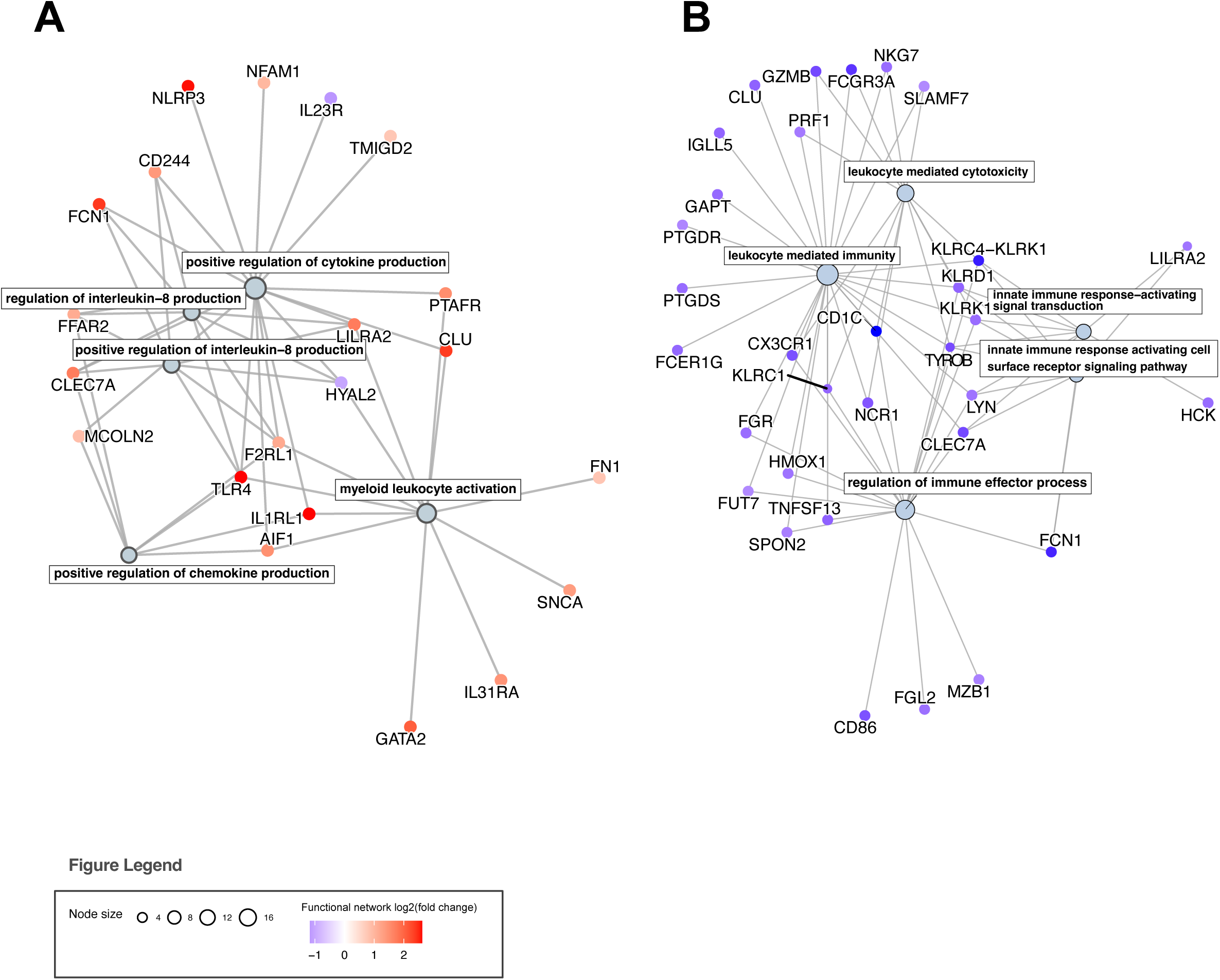
Gene concept network on CD4+ Treg and Teff, Linkages between associated genes and the top 5 enriched GO pathways, with upregulated genes in red and downregulated genes in blue. Teffs: N = 42, Tregs: N = 34. Criteria for inclusion: |log₂FC| ≥ log₂(1.6), p-value ≤ 0.05, FDR ≤ 0.01, and baseMean expression ≥ 8.

### TCR repertoire analysis reveals TRA clonotype expansion in Teff associated with high pain intensity

We further analyzed the T-cell receptor repertoire (TCR) using bulk next generation sequencing (TCR-seq) on the same Teffs and Tregs as those analyzed by transcriptomics. We first compared the diversity of the repertoires between patients with high (N=23, N=22) and low (N=12, N=8) pain by computing the Renyi entropy on normalized TCR-seq data. Renyi entropy curves profile various diversity indices as a function of a parameter alpha (*39*). We found a significantly higher diversity of Teff TRA (T-cell receptor alpha chain) but not TRB (T-cell receptor beta chain) repertoires of OA patients with high pain intensity (*p*<0.001) (**Fig. 5**). No differences were observed in Tregs repertoires. Then our analysis focused on two alpha parameters, the Shannon at alpha=1 which considers both clonotype richness and evenness, and the Berger-Parker index at alpha=Inf which focuses only on the abundance of the most expanded clonotype in the repertoire. Interestingly, we found a significant positive correlation between WOMAC pain score and the Shannon diversity for both TRA (*r* = 0.56, *p*< 0.001) and TRB repertoires (*r* = 0.55, *p*< 0.001). Additionally, we observed a positive correlation between WOMAC pain scores and Berger Parker index in Teff TRA (*r* = 0.58, *p*< 0.001) but not for TRB (*r* = 0.25, *p* = 0.14) repertoires. No significant correlations were observed in Treg cells. These results suggest either bystander activation or recirculation of expanded clones toward the inflammation site in patients with high pain.

**Fig. 5.**
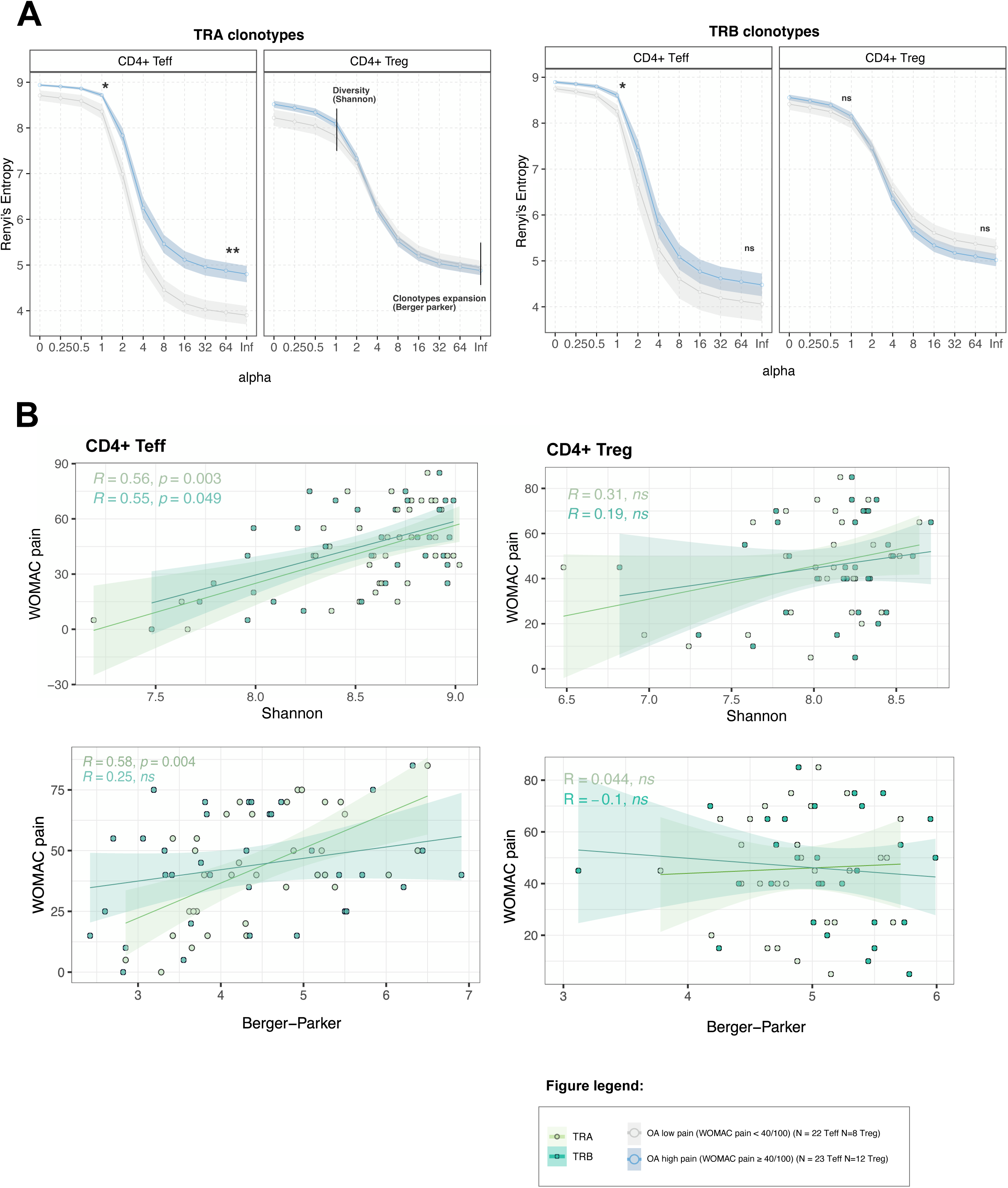
Treg and Teff TCR diversity analysis. A. Renyi’s diversity plots of TRA and TRB amino acid clonotype levels in OA low pain (N=12, N=8) and OA high pain (N = 23, N=22) on CD4+ Teff and Tregs. Mann-Whitney U test was used to assess the difference between groups (p≤0.05). B. Correlation biplot between WOMAC pain and Shannon and Berger-Parker diversity index for CD4+ Tregs and Teff (N = 30, N= 35). Pearson correlation coefficient and statistical p value are plotted for each TCR chain for each diversity index.

### FoxP3+ Treg CTLA4+ subset was decreased with BML and knee effusion - synovitis

To deepen our analysis, we examined previously identified immunological markers associated with OA-related pain and stratified highly symptomatic patients (WOMAC pain ≥40/100) in two groups with either MRI knee effusion-synovitis (N=8) or BML (N=16) allowing to delineate synovial-associated and bone-associated pain phenotype, respectively (Fig. 6A). We found a decrease in Foxp3+ Treg cells expressing CTLA4 in patients with moderate to severe knee effusion-synovitis (N=16) and those with BML (N=6) as compared to patients with low pain intensity (p=0.009, p=0.019). Additionally, effector memory CD8+ T cells (p=0.044), CD95+ Tregs (p=0.049), CD4+ CD45RA-Tregs (p=0.04) percentages were decreased in patients with BML in comparison with low pain intensity (**Fig. 6B-C, S8**). When analyzing serum cytokines, we observed an increase in sTNFR2 levels in patients with BML (N=16) as compared with patients with low pain intensity (WOMAC pain <40/100) (p=0.019) and IL-22 levels were decreased in patients with both knee effusion - synovitis (N=6) and BML (N=16) (p=0.046 and p=0.018) (**Fig. 6D**). Finally, looking at the differential gene expression analysis in Treg and Teff, we observed distinct gene expression signatures that were associated with BML (N=15) and knee effusion - synovitis (N=8). Increased expression of *IL1RL1 and IFITM3* was linked to BML, while *NLRP3, IFI44L*, and *IL31RA* were associated with knee effusion - synovitis (**Fig.7**). Interestingly, *TNFRS11A, SECTM1*, and *KLRC4-KLRK1* were also downregulated in Teffs from patients with BML (N=16), while *TNFSF13, IFI30*, and *NFIL3* were associated with knee effusion - synovitis (N=8).

**Fig. 6.**
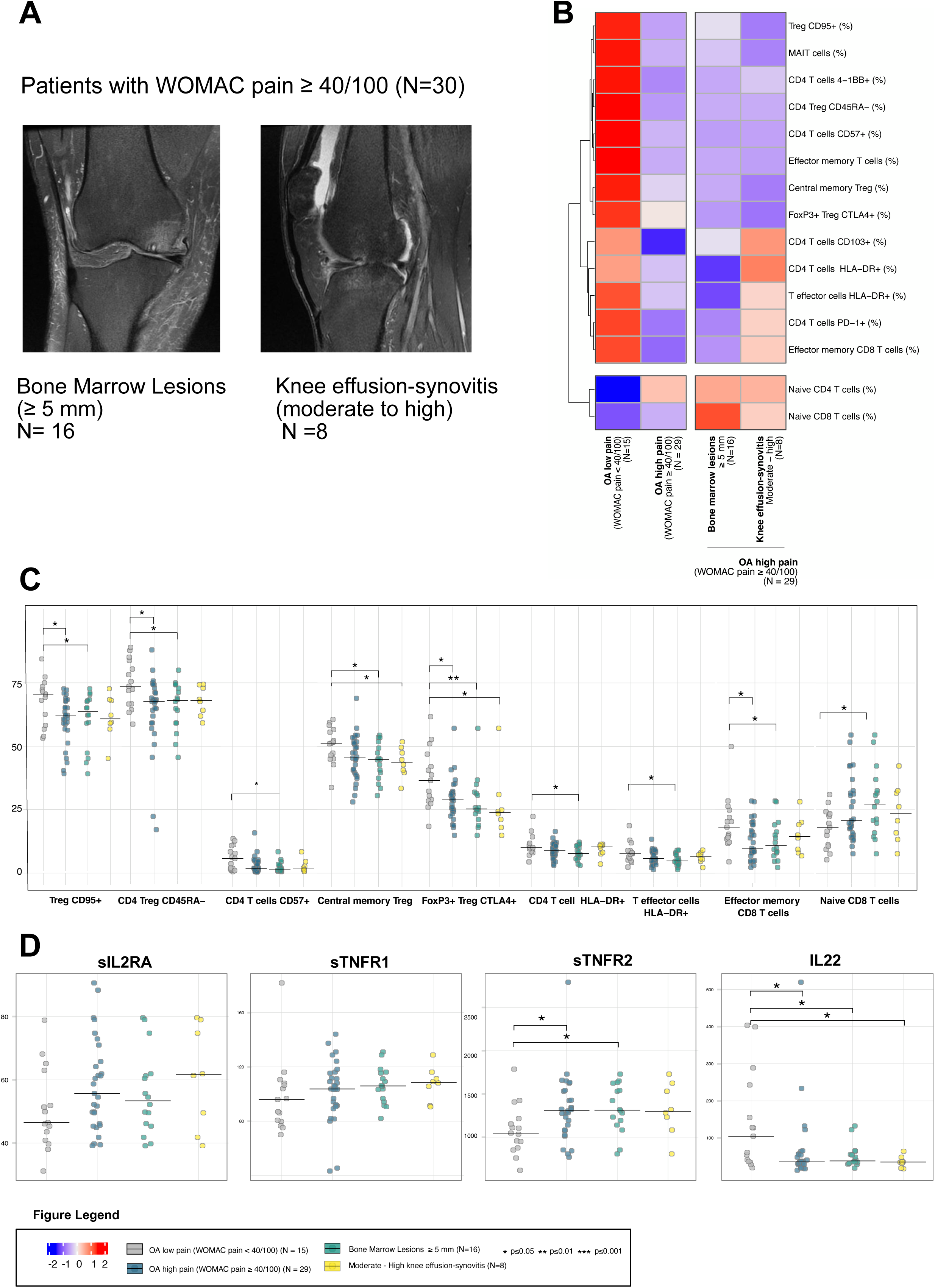
Immunological cell subset and cytokines linked to OA-pain with MRI BML and knee effusion. A. BML and knee representative MRI scan. B. Deep immune phenotyping expression heatmap of the 15 cells subsets associated with WOMAC pain according to low (N=15), high pain intensity (N=29) and BML (N=6) and knee effusion (N=8) represented cell subset mean scaled proportion. C. Scatter plot of cell population percentage (represented on y-axis) specifically associated with high pain intensity, BML or knee effusion compared to low pain intensity group (N=42) using a Mann-whitney U-test (p≤0.05) with patients with low pain intensity as the control group. D. Scatter plot of cytokines MFI-value (y-axis) specifically associated with high pain intensity (N=29), BML (N=16) or knee effusion (N=8) compared to low pain intensity group (N=15) using a Mann-whitney U-test (p≤0.05) with patients with low pain intensity as the control group.

**Fig. 7.**
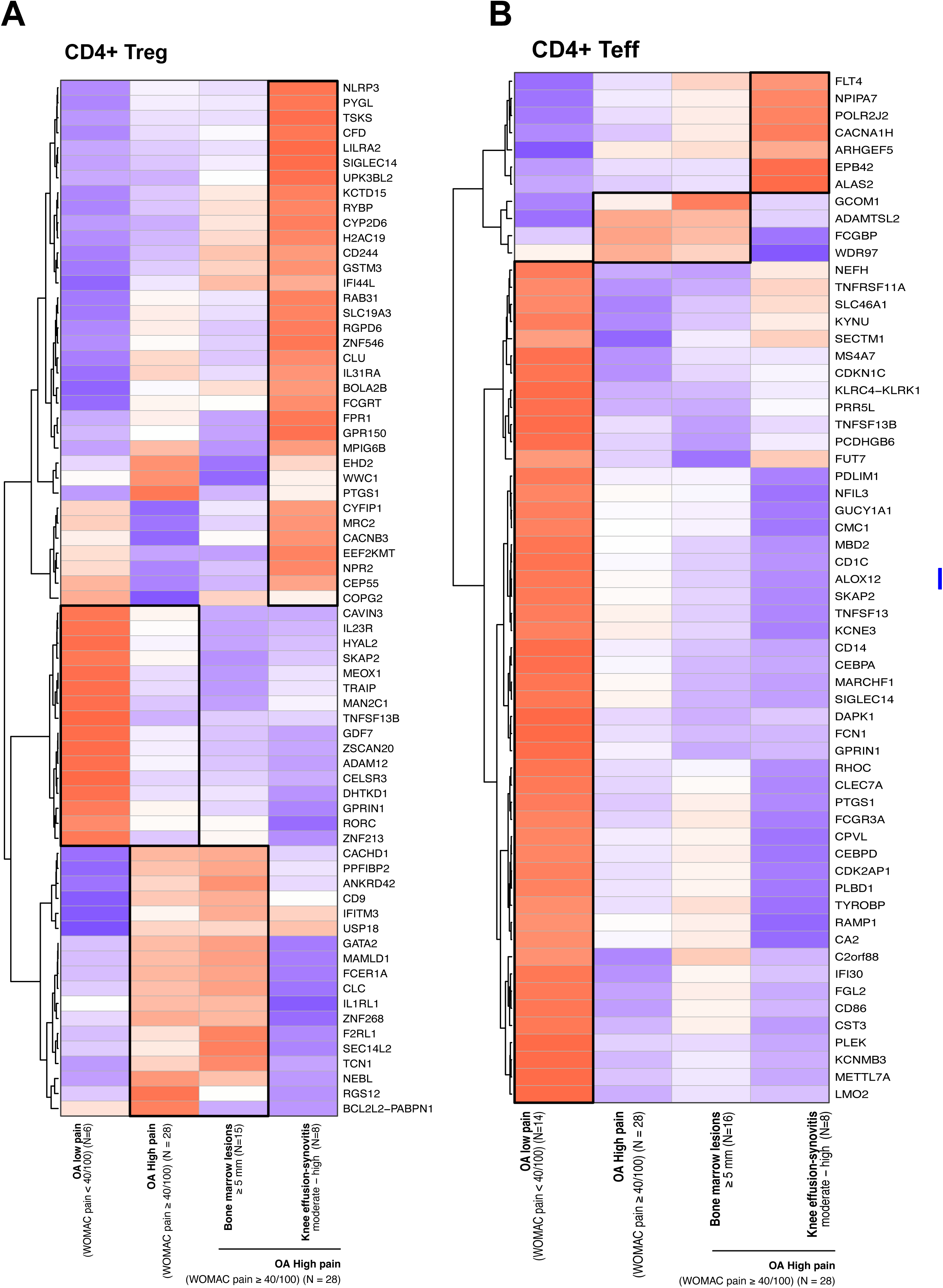
Differential gene expression between low and high OA-related pain with MRI BML and knee effusion - synovitis. A. CD4+ Treg normalized mean-expression heatmap between patients with low pain intensity (N=8), high pain intensity (N=28), BML (N=16), knee effusion - synovitis (N=8). B. CD4+ Teff normalized mean-expression heatmap between patients with low pain intensity (N=14), high pain intensity (N=28), BML (N=16), knee effusion - synovitis (N=8). C. Dot plot of over-representation analysis and enriched GO pathway ordered by gene ratio (N=42) |log2 FC | ≥ log2(1.6), p-value ≤ 0.05, FDR ≤0.01, baseMean expression ≥ 0.8.

## DISCUSSION

In this study, we provide a comprehensive immunological signature associated with OA-related pain using a multi-omic approach and identify a distinct signature associated with WOMAC pain highlighting the potential role of T cells and more particularly Tregs. T cells in OA and OA-related pain have been understudied, although recent results suggest that T cells could emerge as therapeutic targets for chronic pain therapy and Treg could be new key players in the pathophysiology of chronic pain (*40*).

We identified a positive correlation between serum levels of sIL-2RA with WOMAC pain scores. The role of sIL-2RA is not fully understood and whether it acts as a marker of immune regulation or exhaustion remains controversial (*41*). However, elevated levels of sIL-2RA have been associated with autoimmune diseases (*42*, *43*). In vitro models suggest that sIL-2RA may inhibit IL-2 mediated T cell response (*44*). Interestingly low dose IL-2 has been associated with significant arthralgia improvement in a clinical trial across several auto-immune diseases (*45*). In addition our study observed a positive correlation with sTNFR1 and sTNFR2 and WOMAC pain. Increased sTNFR levels have been associated with chronic pain both in peripheral blood and cerebrospinal fluid (CSF) (*46*) and TNFR dual deficient murine model has been associated with chronic pain and inflammation (*47*). Finally, serum IL-22 level was negatively correlated with WOMAC pain. This cytokine has been shown to play a pivotal role in tissue immune regulation and tissue regeneration and has been recently considered as a new therapeutic target for several auto-immune diseases as well as OA (*48*, *49*). Collectively, our cytokine results reinforce the potential role of adaptive immunity and IL-2 in pain modulation in patients with OA.

Deep immunophenotyping revealed a decrease of T cell subsets related to Treg activation and functionality (FoxP3+ CTLA4+ Treg, CD4+ CD57+, Treg CD95+, CD4 Treg CD45RA-). Regulatory T cells have been associated, in experimental studies, with counteracting pain through the inhibition of Th1 responses (*50*), and with inhibition of pain hypersensitivity (*51*). Our previous research has also demonstrated a negative correlation between CD45RA- and CD95+ T cells and inflammatory states as well as auto-inflammatory diseases (*52*).

Strikingly, differential gene expression analysis of Tregs between patients with low and high WOMAC pain scores identified an upregulation of inflammasome-related genes suggesting overrepresentation pathways related to innate immunity suggesting a pro-inflammatory state of Tregs. Van der Veeken et al. have demonstrated that inflammation-experienced Treg reversed many activation-induced changes and could gradually lose its enhanced suppressive function over time (*53*). Altogether our results suggest a potential Treg dysfunction associated with OA-related pain. Additionally, Teff transcriptomic analysis showed an upregulation of genes associated with bone remodeling such as *RANK* and a downregulation of genes associated with NK and CD8+ responses, and an over-representation of pathways related to cell cytotoxicity. Activated NK and T cells have been described to move into the synovium in OA and RA (*54–56*) and several studies have underscored a divergence between synovium and peripheral blood transcriptomic profile (*57*, *58*). Additionally, other immune cells have been observed migrating into the dorsal root ganglion (DRG) and sensory neurons, playing a role in OA-related pain mechanisms (*59*). Considering the Shannon index as a measure of diversity, independent of species abundance, the TRA and TRB repertoires of Teff cells are more diverse in OA patients with high pain. However, at the level of large clonotype expansions, no significant differences are observed in the TRB repertoire, whereas the TRA repertoire is notably more diverse. This could be explained by increased activation of dual-alpha receptor T cells in high pain, which share similar TRB expansions with the low pain group but exhibit a more diverse TRA repertoire. This diversity may result from bystander activation or the recirculation of expanded clones to the site of inflammation in patients experiencing high pain.

MRI-based stratification found a decrease of specific cell subsets, such as FoxP3+ Treg and CTLA4+ cells, associated with BML and knee effusion-synovitis, amd offers valuable insights into OA-related pain phenotypes that are bone-associated or synovial-associated pain phenotypes (*60*).

Although the sample size of our study could limit our ability to fully characterize pain phenotypes or endotypes, these findings highlight the potential of targeted approaches. In addition, our study did not address central sensitization, a critical factor in pain phenotyping. Accurately defining pain phenotypes in OA remains a significant challenge but is essential for advancing precision medicine in this field (*21*, *61*). The limited sample size of our study restricted the use of adjustments for multiple testing. While this analysis is exploratory, further studies with larger cohorts are required to validate our findings. In addition, analysis of synovial tissue alongside matched peripheral blood are needed to better understand Teff-Treg modulation and possible role in contributing to pain in patients with OA. The use of synovial biopsies could provide new insights and perspectives (*62*) and in vitro and experimental models could help us enhance our comprehension of the underlying pathophysiology (*63*). Our study provides a detailed exploration of the immunological landscape associated with OA-related pain. Using a multi-omic approach that combines cytokine profiling, deep immunophenotyping, transcriptomics of sorted Treg and Teff cells, and TCR repertoire analysis for each patient, we identified a pro-inflammatory signature in Tregs associated with WOMAC pain, suggesting new pathogenic hypotheses on the roles of T cells and Tregs in pain. These findings open new avenues for the evaluation of therapeutic strategies that target T cells to manage chronic pain in OA patients.

## MATERIAL AND METHODS

### Study Design

The TRANSIMMUNOM study (NCT02466217) is a multi-center clinical trial aiming to cross-phenotype using a multi-omic approach patients with autoimmune and/or autoinflammatory diseases, based on the continuum from auto-inflammatory to auto-immune diseases, delineated by McGonagle and McDermott (*64*). TRANSIMMUNOM included patients with knee OA classified as a low-grade autoinflammatory disease within this continuum. To ensure standardized data collection, we utilized electronic case-report forms (E-CRF) to record clinical, biological, and radiological data for each patient. The comprehensive methodology and procedures for this study are detailed in the Transimmunom protocol, as described by Lorenzon et al (*32*). In addition, 135 mL of peripheral blood were collected from each participant in the study. From October 2017 to June 2020, 46 patients with knee OA fulfilling ACR criteria were recruited from the rheumatology department of Saint-Antoine Hospital in Paris, France (*33*). To be included in the study, patients had to have either unilateral or bilateral radiographic knee OA with a KL score of 2 or 3 on the most affected compartment of the target knee, in order to exclude late-stage OA and have homogenous groups in terms of radiographic severity, and be at least 35 years old. Patients were excluded if they had a total knee replacement, a KL score of 1 or 4, a co-existing fibromyalgia, inflammatory rheumatic disease, crystal induced arthropathies, secondary OA related to rare genetic disorder or avascular necrosis of the knee. The inclusion and exclusion criteria are presented in **table S1**. There was no cut-off in terms of pain intensity in order to include a wide range of pain symptom severity. The TRANSIMMUNOM study received approval from the institutional review board of Pitié-Salpêtrière Hospital (CPP Ile de France VI) on 16 July 2015. The study adhered to the principles outlined in the Declaration of Helsinki and followed guidelines for good clinical practice. All participants provided written informed consent prior to enrollment in the study.

### Clinical assessment

Patients underwent a standardized clinical questionnaire, clinical exam, blood sample for research and routine biology. Clinical variables included: demographic variables such as age, gender, body mass index (BMI), menopause, and knee OA disease duration and high sensitive C-reactive protein (CRP) assessment. Pain and symptoms scores were collected using the Western Ontario and McMaster Universities Arthritis Index (WOMAC) pain, function, stiffness subscores (*65*), Hospital anxiety and depression score (HADS) (*66*), *Douleur neuropathique 4* questionnaire (DN4) to assess neuropathic-like pain component (*67*). Patients were also divided into low and high pain intensity groups using a WOMAC pain threshold ≥ 40/100 corresponding to the patient acceptable symptom state (PASS) (*36*). Information related to treatment including acetaminophen, oral and local non-steroidal anti-inflammatory drugs (NSAIDs), and weak opioids (defined as the intake of codeine, dihydrocodeine, or tramadol) were also collected.

### Knee joints X-rays and MRI semi-quantification

Knee X-ray included weight bearing antero-posterior, lateral, Schuss and femoro-patellar defile and quantified using the Kellgren Lawrence score (*34*). Non injected MRIs were performed on the most symptomatic knee using a standardized protocol using a 1.5 MRI with a standardized protocol using a multi-planar weighted T1 sequence and intermediate proton density (fat-sat). Semi-quantitative scoring of joint effusion and subchondral changes was conducted by a senior musculoskeletal radiologist (LM) who was blinded to clinical, biological, and radiographic data, using the effusion grades in the tibiofemoral joint (TFJ) and patellofemoral joint (PFJ) were combined and scored as follows: 0 = absent, 1 = small, 2 = moderate, and 3 = severe. knee effusion - synovitis was characterized by a score of ≥ 2, indicating moderate or severe joint effusion. Subchondral edema was graded as follows: 0 = absent, 1 = minimal (<5 mm diameter), 2 = moderate (5 mm to 2 cm diameter), and 3 = severe (>2 cm diameter). Subchondral edema was defined with a grade of ≥ 2, corresponding to a diameter of ≥ 5 mm.

### Serum cytokines

Serum cytokines, chemokines and soluble receptors were quantified by Luminex® multiplex technology. Five MILLIPLEX® kits were used: Human Cytokine/Chemokine, Human High Sensitivity T Cell, Human Th17, Human TGFbeta1 and Human Soluble Cytokine Receptor Magnetic Bead Panel 96-well plate assay (Merck). Data acquisitions were performed on a Luminex® MAGPIX instrument and data were analyzed using the xPONENT® software. Calibration and verification of the instrument were performed one week before each experiment. For each kit and each batch performed, two technical controls, with expected range, were duplicated as quality control. During analysis, the standards with percentage of recovery <80 or >120 were removed. Mean fluorescence intensity data using a 5-parameter logistic or spline curve-fitting method was used for calculating analyte concentrations in samples (based on standard concentrations). Cytokines, chemokines and soluble receptors quantified are summarized in **table S6**

### Flow cytometry and deep immunophenotyping

Cytometry profiling of patient blood samples was performed using 12 flow cytometry panels developed for the TRANSIMMUNOM study (*37*, *38*). As previously described, Duraclone technology was used in whole blood, which was preferred over PBMC to reduce technical manipulations and minimize loss or reduction of phenotype. In detail, five panels were designed to analyze B cells, natural killer (NK) cells, monocytes, dendritic cells, mucosal-associated invariant T (MAIT) cells and myeloid-derived suppressor cells. Four panels were designed to investigate T-cells activation, migration, memory and polarization phenotypes and two panels were focused on regulatory T cell populations. An extra panel was also developed to identify main immune cell populations and included numeration beads allowing to determine the absolute counts of all immune cell populations. All acquisitions were performed on a Gallios cytometer and flow cytometry data were analyzed with Kaluza software version 1.3 (Beckman Coulter). For each cell population percentages of parent population, absolute count of cells and mean fluorescence intensities were obtained. Panel composition, cell populations and gating strategies are resumed in **Fig. S2**, **table S2**.

### T cell sorting

TCR-dequencing (TCR-Seq) and RNA sequencing data generated from sorted T-cell subsets from peripheral blood samples (30mL). PBMC were isolated using a Ficoll-Paque density gradient. CD4+ T cells were enriched from the PBMCs using EasySepTM magnetic beads, followed by separation into Treg (CD25+CD127loCD4+) and Teff (CD25-CD127+CD4+) subsets according to manufacturer’s recommendations. The purity of these cell subsets was verified by flow cytometry using CD4 and FoxP3 antibodies using a threshold ≥ 80%. The sorting accuracy was ranging from 92% to 98% for Teffs and was ranging from 85% to 98% for Tregs.

### Treg and Teff Transcriptomic analysis

Total RNA was extracted from sorted Treg and Teff cells using the RNAqueousTM – Micro Total RNA Isolation Ki Kit (Invintrogen Waltham, MA) following the manufacturer’s instructions. The isolated RNA was reverse-transcribed with an Oligo(dT) primer and cDNA amplification was amplified, fragmented and tagged using the SMARTer® Ultra® Low Input RNA Kit for Sequencing—v4 (Takara Bio USA, IncMountain View, CA) Quality control was performed using either the Bioanalyzer 2100 (Agilent) or the Tapestation 4200 (Agilent) on 10 ng of RNA for Tregs and 100 ng for Teffs. RNA Integrity Number (RIN) was performed to asset extraction quality (RIN threshold ≥ 7). RNA-seq profiling was performed on Illumina Next-Seq 500 (75bp, paired-end) to reach on average 15M reads per sample. Sequenced reads were trimmed using Trim Galore, and aligned using Salmon on the release GRCCh38.p13 of Ensembl Human Transcriptome reference (*68*, *69*). For the transcriptomics analysis, quality controls were performed at both sample and gene levels. Genes with counts below 20 in fewer than 10 patients for Teff cells and 5 patients for Tregs were filtered out from the normalized matrix. Quality control measures are summarized in **Fig. S6-7**. Differential gene expression analysis was conducted using DESeq2 (version 1.38.3) (*70*) with a log2 fold change threshold of ≥ |log2(1.6)|, p ≤ 0.01, and a baseMean expression ≥ 8. Functional analysis included over-representation analysis with Gene Ontology pathway analysis with (p-value ≤ 0.5, log2 fold change ≥ |1.6|, baseMean expression ≥ 0.8, false discovery rate (FDR) q-value ≤ 0.05) (*71*). Gene concept network was performed using the enrichplot, DOSE, clusterProfiler package (*72*, *73*).

### TCR libraries preparation, sequencing, data preprocessing and quality control

Total RNA was extracted and quantified using Bioanalyzer 2100 (Agilent) or Tapestation 4200 (Agilent). 10 ng for Tregs and 100 ng for Teffs of RNA were used for TCR transcripts amplification using SMARTer Human TCR a/b Profiling Kit v1 (Takarabio) (*13*). TCR cDNA was then sequenced by Next Generation Sequencing (NGS) using HiSeq 2500 (Illumina) and NOVAseq 6000 by the EquiPex LIGAN core facility (Lille, France) or by IGenSeq at the Paris Brain institute (Paris, France). Quality control (QC) of the raw data was performed using FastQC and multiQC (*74*). Raw data were then aligned using MiXCR version 3.0.13 (*75*). When samples were sequenced with the NOVA-seq 6000, the DeconTCR package, developed by the i3 lab, was used to correct index hopping contamination. Cured TCR aligned data underwent quality control using the QtCR package. Prior to TCR-seq data analysis, singletons, unproductive sequences and sequences beyond CDR3 length limits were filtered out by AnalyzAIRR package. To limit the impact of the number of cells differing from one sample to another, a normalization was applied by downsampling the total number of sequences in each sample to 10,000 using AnalyzAIRR package.

### Statistical Analysis

Clinical variables were reported as mean ± standard deviation and stratified into groups based on low and high pain intensity. Differences between the low and high pain intensity groups were assessed using Student-test for continuous variables and chi-square tests for categorical variables. Flow cytometry and cytokine analyses employed unsupervised non-parametric approaches, specifically Spearman correlation coefficient using an absolute correlation threshold |r| >0.2 and p-value threshold ≤ 0.05. Differential gene expression was assessed using DESeq2 with negative binomial distribution to model the count data, accounting for overdispersion, and applied the Wald test to assess the significance of differential expression between conditions by testing whether the estimated log fold changes differ significantly from zero, DESEq2 and Salmon packages (*69*, *70*). Functional analysis was conducted with clusterProfiler package (*73*). TCR-Seq analysis utilized Renyi’s entropy for entropy calculation and Shannon and Berger-Parker diversity indices for diversity assessment. Analysis was conducted using DeconTCR and AnalyzAIRR packages (*76*) and TRA-TRB correlation analyses with diversity indices were based on the Pearson coefficient. Subset analysis was conducted on patients with high pain (WOMAC pain ≥ 40/100) and moderate to severe knee effusion-synovitis (N=8) or BML) on MRI (N=16). A Mann-Whitney test was performed for statistical comparisons. Given the exploratory nature of this study and the limited patient cohort size, adjustments for multiple testing were not applied (*77*, *78*).

## Acknowledgements

We would like to thank the patients who participated in the Transimmunom study, as well as the administrative and medical teams of the I3 Lab and the Clinical Research Center of Sorbonne University. We extend our thanks to the investigators involved in patient recruitment from the Rheumatology Department of Saint-Antoine Hospital (AP-HP), including Dr. Camille Deprouw, Dr. Sandra Desouches, Dr. Ariane Do, Dr. Karine Louati, Dr. Sabine Trellu, Dr. Juliette-Louise Petit, Dr Julien Champey and Dr. Catherine Beauvais.

## Author’s contribution

RL, CR, SH, AC, CA, FB, and JS recruited patients for the study. JD, FP, AR, MBa, and MR performed sample collection, flow cytometry, and cytokine analysis. LM conducted radiographic severity assessment and MRI analysis. MB, PS, HV, LA, VM, NC, GF, VD, PB, and MR performed cell sorting, RNA extraction, and TCR sequencing. YM and NT performed RNA sequencing. MB, PS, JS, NT, and MR conducted quality controls and pre-processing. MB, PS, and MMS performed TCR analysis. MB conducted flow cytometry, cytokine, mRNA, and functional analysis. SB, AJB, and AS provided essential feedback. PS and NT provided feedback regarding the analyses. CA was involved in project administration. FB, DK, EMF, and JS conceptualized the study. All authors revised the manuscript.

## Data and materials availability

Upon manuscript acceptance, the data and code will be made publicly accessible on GO and GitHub. Supporting data values for figures are also included in supplemental files.

## Supplementary

Supplementary Figures 1 - 9

Supplementary Tables 1-9

## Abbreviation list

BMI: Body Mass Index
BML: Bone Marrow Lesions
CCR: Chemokine receptor
CD: Cluster of differentiation
CTLA-4: Cytotoxic T-lymphocyte–associated antigen 4
DC: Dendritic cells
DN4: Douleur neuropathique 4
FoxP3: Forkhead box P3
GCSF: Granulocyte Colony-Stimulating Factor
GITR: Glucocorticoid-Induced TNFR-Related
HADS: Hospital Anxiety and Depression Scale
HLA-DR: Human leukocyte antigen-DR isotype
hsCRP: High sensitive C-reactive protein
ICOS: Inducible costimulator
IFN: Interferon
ILC: Innate lymphoid cells
Ig: Immunoglobulin
IP16: Interferon-Inducible Protein 16
KL: Kellgren Lawrence
LAG-3: Lymphocyte activation gene 3
LAP: LC3-associated phagocytosis
LFT: Lateral femoro-tibial
MAIT: Mucosal-associated invariant T cells
MCP: Monocyte Chemoattractant Protein
MFT: Medial femoro-tibial
MRI: Magnetic Resonance Imaging
NK: Natural Killer
NKG2D: Natural Killer group 2 member D
NSAIDs: Non-steroidal anti-inflammatory drugs
OA: Osteoarthritis
PD-1: Programmed Cell Death Protein 1
sIL1RL1: Soluble Interleukin-1 Receptor-Like 1
sIL2RA: Soluble Interleukin-2 Receptor Alpha
sRAGE: Soluble Receptor for Advanced Glycation End-products
sTFNR: Soluble Tumor Necrosis Factor Receptor
TEMRA: Terminal effector memory T cells
TNF: Tumor Necrosis Factor
TCR: T cell receptor
Treg: T regulatory cell
WOMAC: Western Ontario and McMaster Universities Arthritis Index.

