## Supplements for "Immunological Profiling in Knee Osteoarthritis: Treg Dysfunction as Key Driver of Pain"

**SUPPLEMENTARY:**

**Figures Legend:**

**Fig. S1 | Spearman Correlation plot between WOMAC pain and cytokines presented in MFI.** Only Cytokines significantly associated with WOMAC pain were shown, p ≤ 0.05, |r| ≥0.02.

**Fig. S2 | Deep immunophenotyping gating strategy adapted from Pitoiset et al.** Panel and targeted cell populations were highlighted in color. Cell populations involved in the numeration panel allowing for beads counts and identification and quantification of all populations were highlighted in blue.

**Fig. S3 | Spearman Correlation plot between WOMAC pain cell populations, presented as percentage**. Only cell populations significantly associated with WOMAC pain were shown, p ≤ 0.05, |r| ≥0.02.

**Fig. S4 | Flow cytometry analysis of immune cell subsets associated with OA-related pain in patients matched for age, sex, and BMI, comparing those with low and high pain intensity.** Treg function panel including CD45RA- subpopulation, Treg phenotype panel including CTLA+ FoxP3+ subpopulation T naive/memory panel, including Central Memory Tregs (CD95+ Treg) Effector Memory CD8+ T cells.

**Fig. S5 | MSD - VSD CD4+Teff (N = 42) and for CD4+ Treg (N = 34**). Samples projection according to low and high pain intensity status.

**Fig. S6 | Treg mRNA sequencing quality control (N=30).** A. Dispersion plot. B. Treg samples Cook’s distance. C. MDS-VSD Treg projection according to batch sex BMI and age.

**Fig. S7 | Treff mRNA sequencing quality control (N=42).** A Dispersion plot. B. Treg samples Cook’s distance. C. MDS-VSD Treg projection according to batch sex BMI and age.

**Fig. S8 | Scatter plot and Wilcoxon analysis between low pain intensity group and pain phenotypes** (high pain intensity, BML and knee effusion). p ≤ 0.05, |r| ≥0.02.

**Tables legend:**

**Table S1 | Inclusion and exclusion criteria for OA patients in the Transimmunom study.**

**Table S2 | Cytokines and Panel measured in the Transimmunom study.** Cytokines were measured using four Luminex panels including Cytokines-Chemokines kit (CC-CK), Highly sensitive kit (HS), Receptor kit, and TH17 kit. Measurements were expressed in MFI (mean fluorescence intensity) and included 62 cytokines.

**Table S3 | Spearman correlation of cytokines significantly associated with WOMAC pain and DN4.** Significant correlations between cell subsets and symptoms were represented using a threshold of p ≤ 0.05 and |r| ≥ 0.2.

**Table S4 | Deep immunophenotyping cells populations and panels.** Cell populations were measured across 12 panels, including a general panel, B cell panel, NK-NCR cell panel, T cell activation and naive/memory T cell panel, MAIT-NKT cell panel, T cell migration panel, CD4+ T cell polarization panel, T cell activation panel, Treg function panel, DC/monocyte panel, Treg phenotype panel, and Myeloid-ILCs panel. The measurements were expressed in percentage (%) and include 269 cell populations across these panels.

**Table S5 | Spearman correlation of cell populations significantly associated with WOMAC pain and DN4.** Significant correlations between cell subsets and symptoms were represented using a threshold of p ≤ 0.05 and |r| ≥ 0.2.

**Table S6 | Differentially expressed genes in Treg of patients with low (N=6) and high pain intensity in Treg (N=28).** (|log2 FC | ≥ log2(1.6), p-value ≤ 0.05, baseMean expression ≥ 8).

**Table S7 | Differentially expressed genes in Teff of patients with low (N=14) and high pain intensity in Treg (N=28)** (|log2 FC| ≥ log2(1.6),, p-value ≤ 0.05, baseMean expression ≥ 8.

**Table S8 | Non parametric Mann-Whitney analysis comparing cell subsets and patients with WOMAC pain score ≥40/100 and different pain phenotypes (neuropathic pain (N=10), BML (N=16), knee effusion (N=16)).** For each pain phenotype variable, comparisons were made with groups without neuropathic pain (N=42), without subchondral edema and pain (N=23), and without knee effusion (N=31). A significant p-value was defined with a threshold of ≤ 0.05.

**Table S9 | Non-parametric Mann-Whitney analysis comparing cytokines and patients with WOMAC pain score ≥40/100 and different pain phenotypes (neuropathic pain (N=10), BML (N=16), knee effusion (N=16)).** For each pain phenotype variable, comparisons were made with groups without neuropathic pain (N=42), without subchondral edema and pain (N=23), and without knee effusion (N=31). A significant p-value was defined with a threshold of ≤ 0.05.

**Fig. S1 | Spearman Correlation plot between WOMAC pain and cytokines presented in MFI**. Only Cytokines significantly associated with WOMAC pain were shown, p ≤ 0.05, |r| ≥0.02.

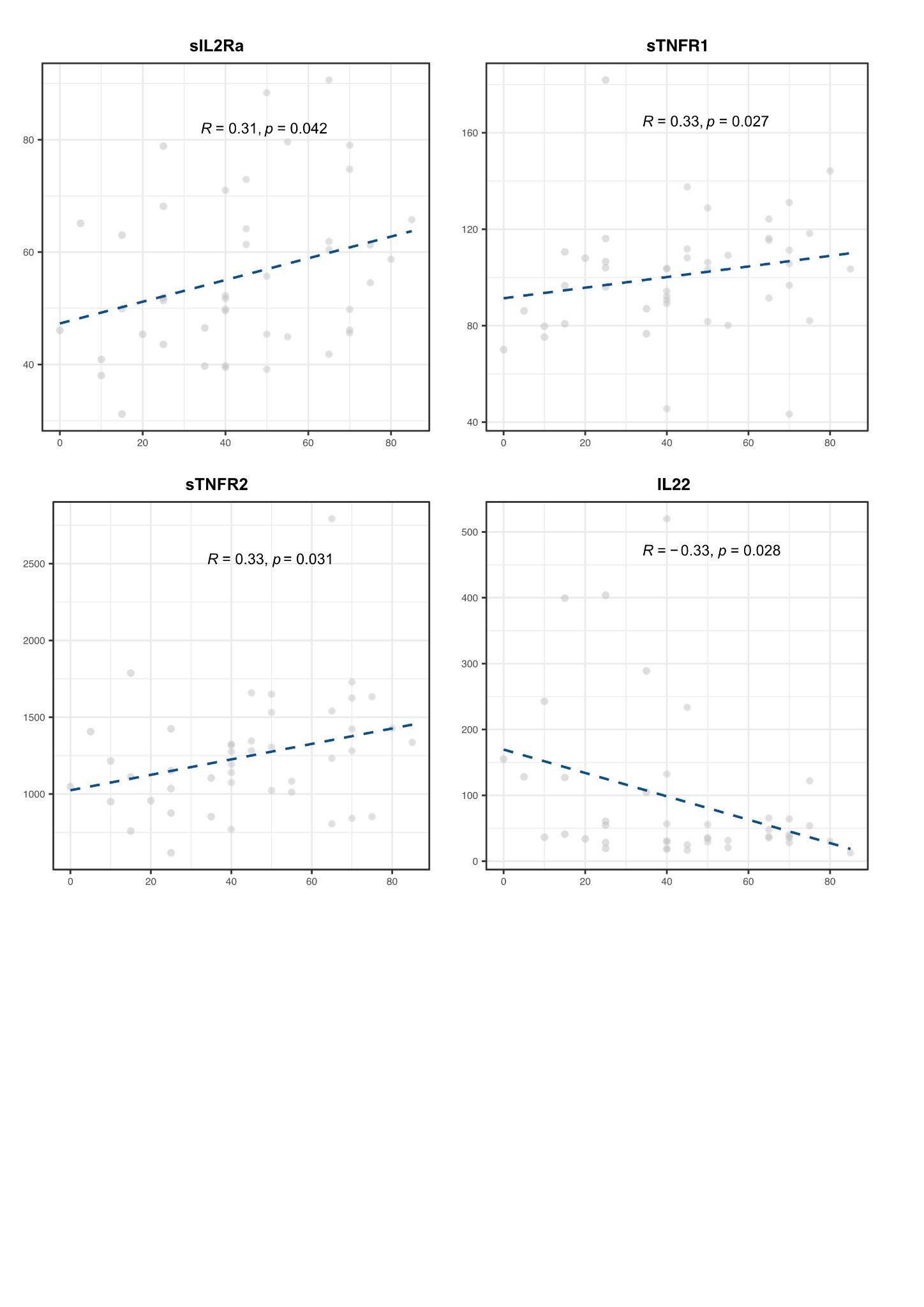

IL, Interleukin, sTNFR, soluble Tumor Necrosis Factor Receptor

**Fig. S2 | Deep immunophenotyping gating strategy adapted from Pitoiset et al.** Panel and targeted cell populations were highlighted in color. Cell populations involved in the numeration panel allowing for beads counts and identification and quantification of all populations were highlighted in blue.

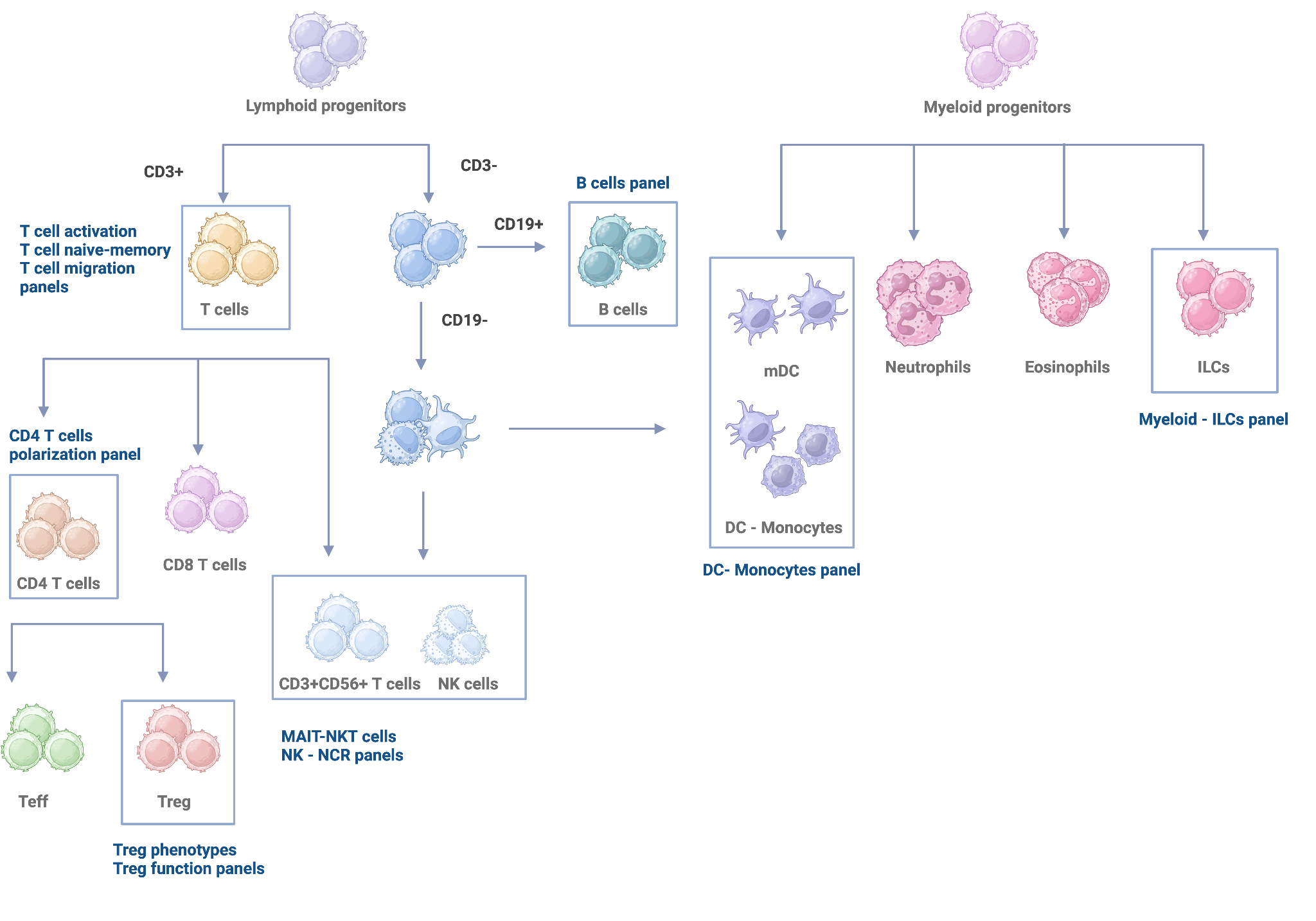

CD, Cluster of differentiation; ILC, Innate lymphoid cells; MAIT, Mucosal-associated invariant T cells; NK, Natural Killer, ILC, Innate lymphoid cells; NK, Natural killer; NKT, Natural Killer T cells; Teff T effectory; Treg, T regulatory

**Fig. S3 | Spearman Correlation plot between WOMAC pain cell populations, presented as percentage.** Only cell populations significantly associated with WOMAC pain were shown, p ≤ 0.05, |r| ≥0.02.

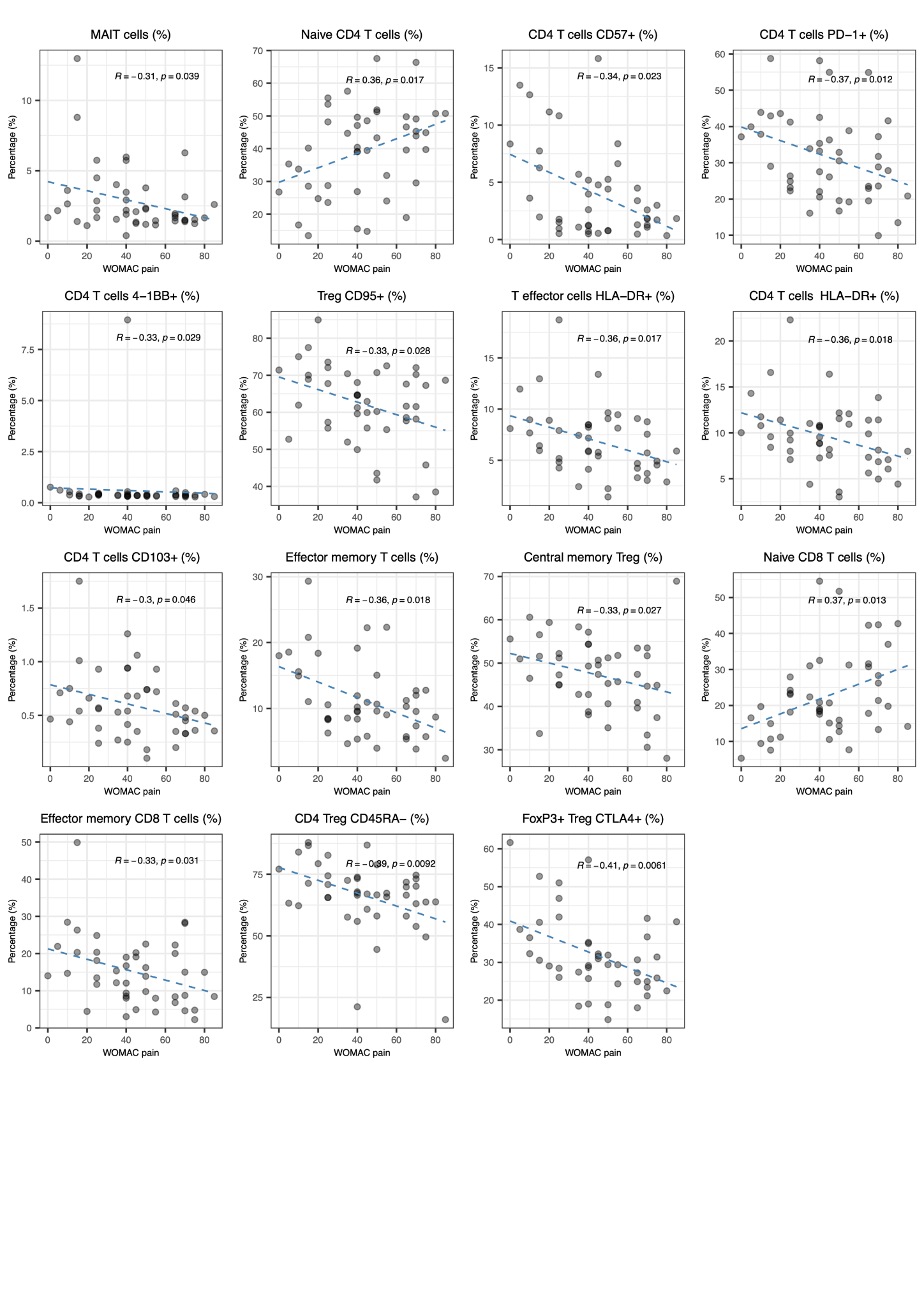

CD, Cluster of differentiation; HLA-DR, Human leukocyte antigen-DR isotype; MAIT, Mucosal-associated invariant T; PD-1, Programmed Cell Death Protein 1; WOMAC, Western Ontario and McMaster universities arthritis index

**FIg S4 | Flow cytometry analysis of immune cell subsets associated with OA-related pain in patients matched for age, sex, and BMI, comparing those with low and high pain intensity.** Treg function panel including CD45RA- subpopulation, Treg phenotype panel including CTLA+FoxP3+ subpopulation T naive/memory panel, including Central Memory Tregs (CD95+ Treg) Effector Memory CD8+ T cells

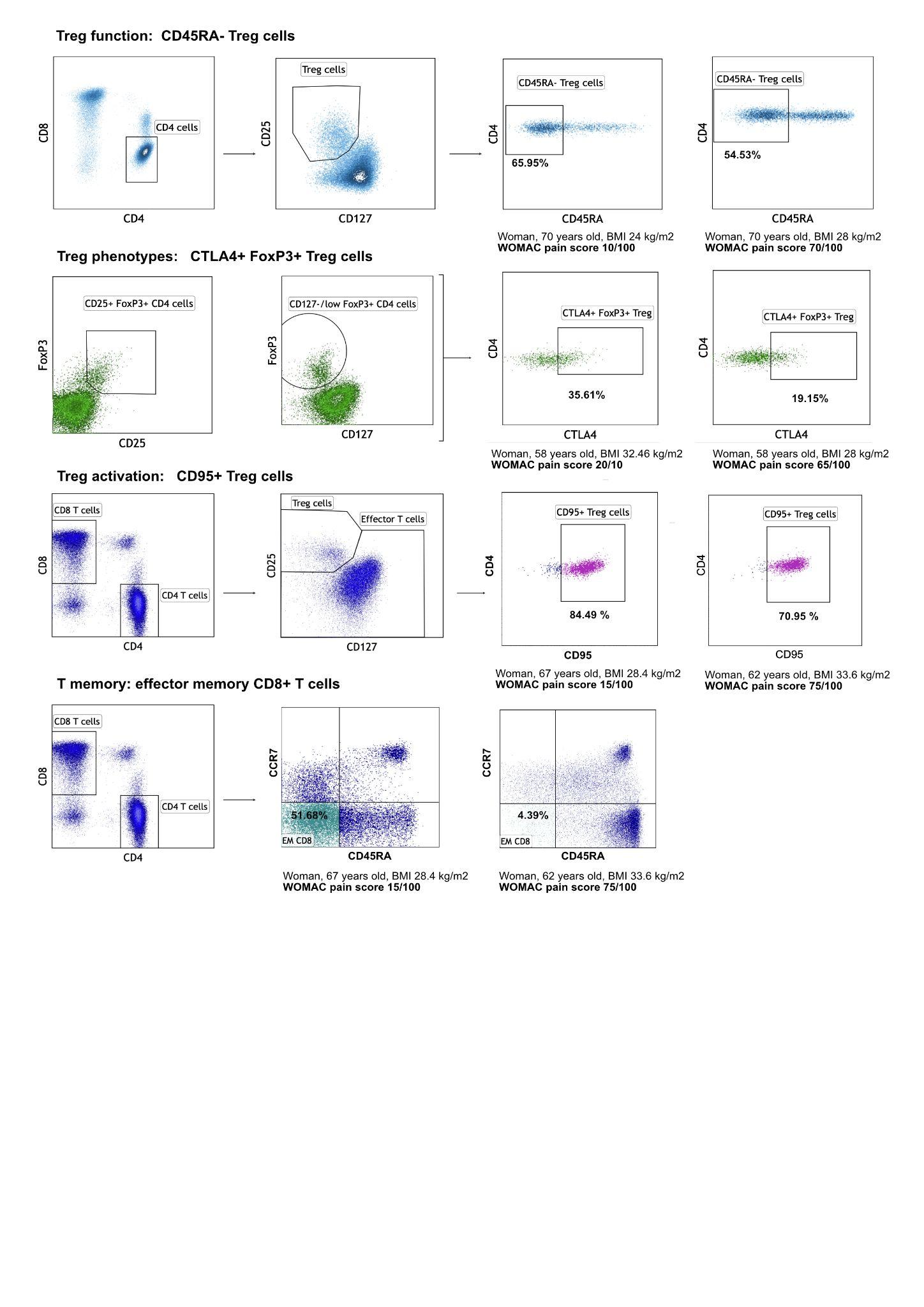

**Fig. S5 | MSD - VSD CD4+ Teff (N = 42) and for CD4+ Treg (N = 34**). Sample projection according to low and high pain intensity status.

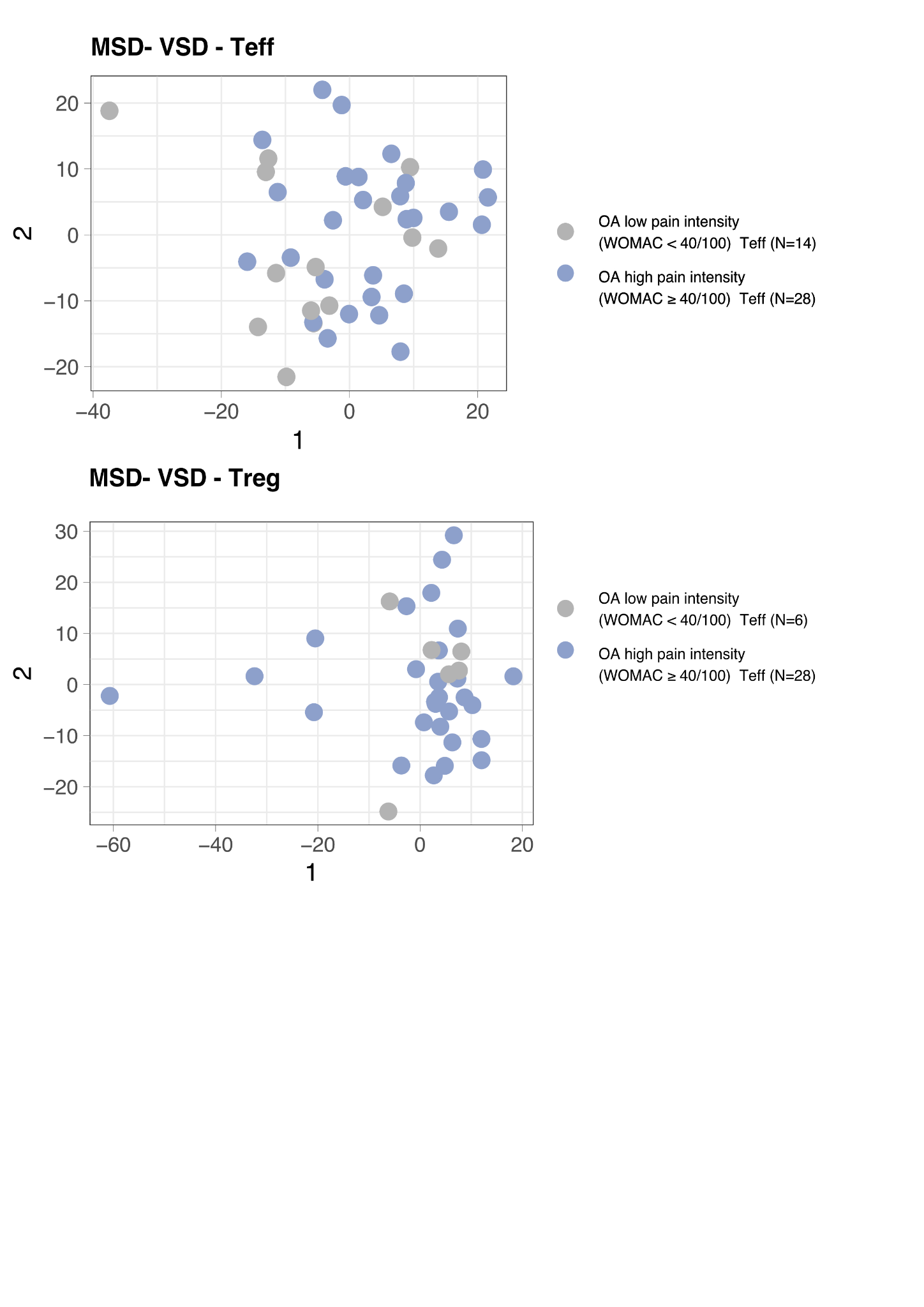

**Fig. S6 | Treg mRNA sequencing quality control** (N=30). A Dispersion plot. B. Treg samples Cook’s distance. C. MDS-VSD Treg projection according to batch sex BMI and age.

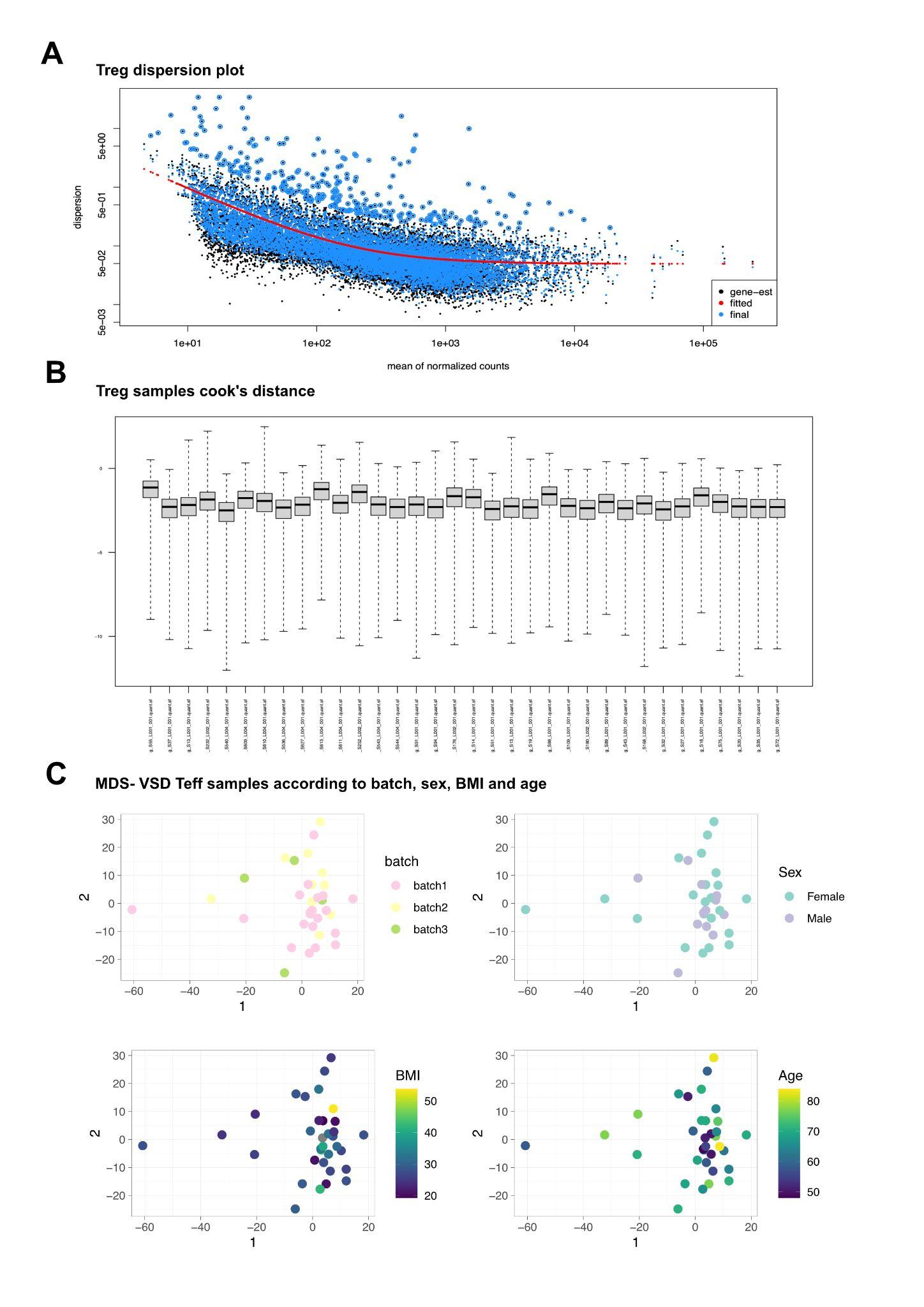

**Fig. S7 | Teff mRNA sequencing quality control** (N=42). A Dispersion plot. B. Treg samples Cook’s distance. C. MDS-VSD Treg projection according to batch sex BMI and age.

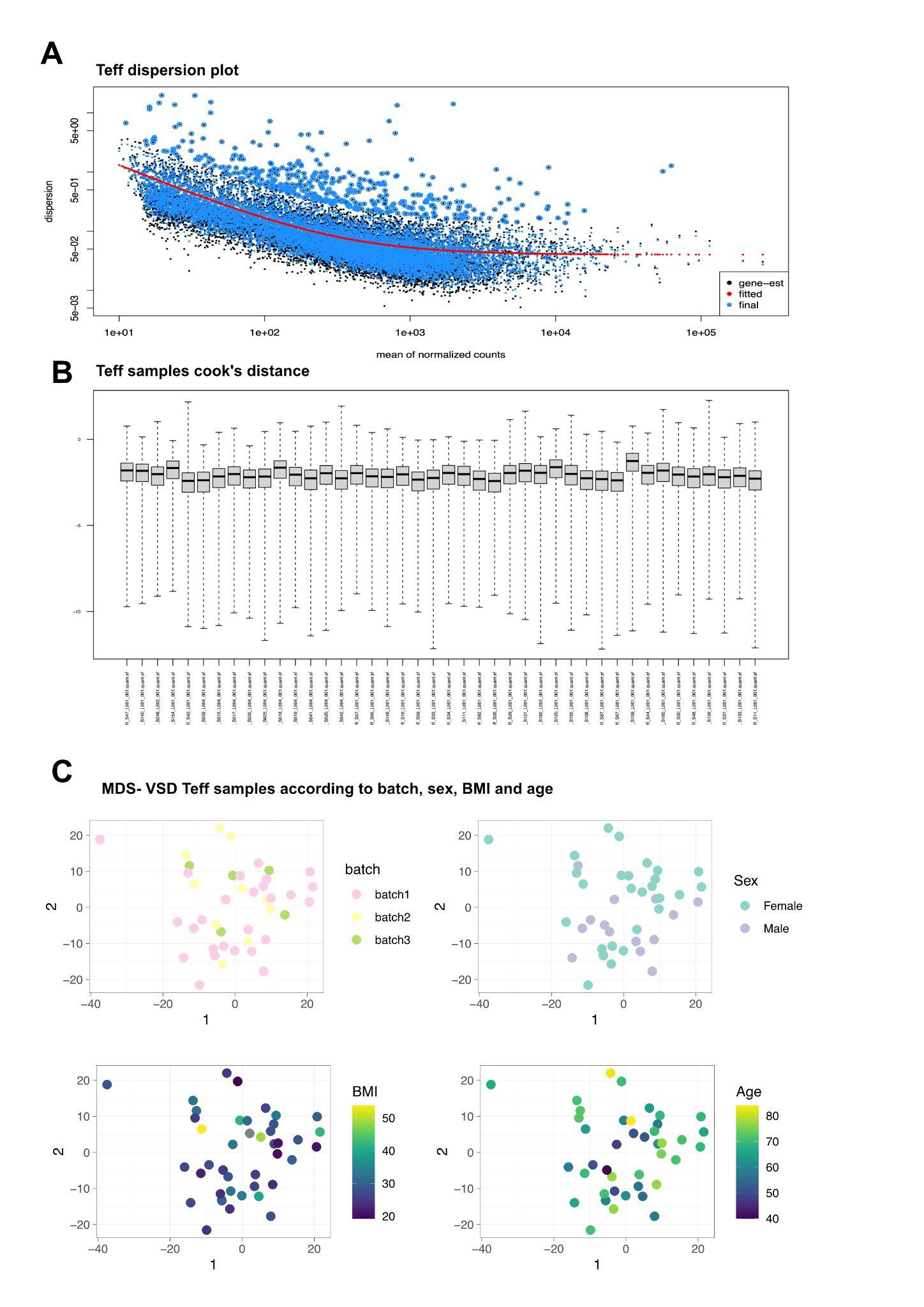

**Fig. S8 | Scatter plot and Mann-Whitney analysis between low pain intensity group and pain phenotypes** (high pain intensity, BML and knee effusion). p ≤ 0.05, |r| ≥0.02.

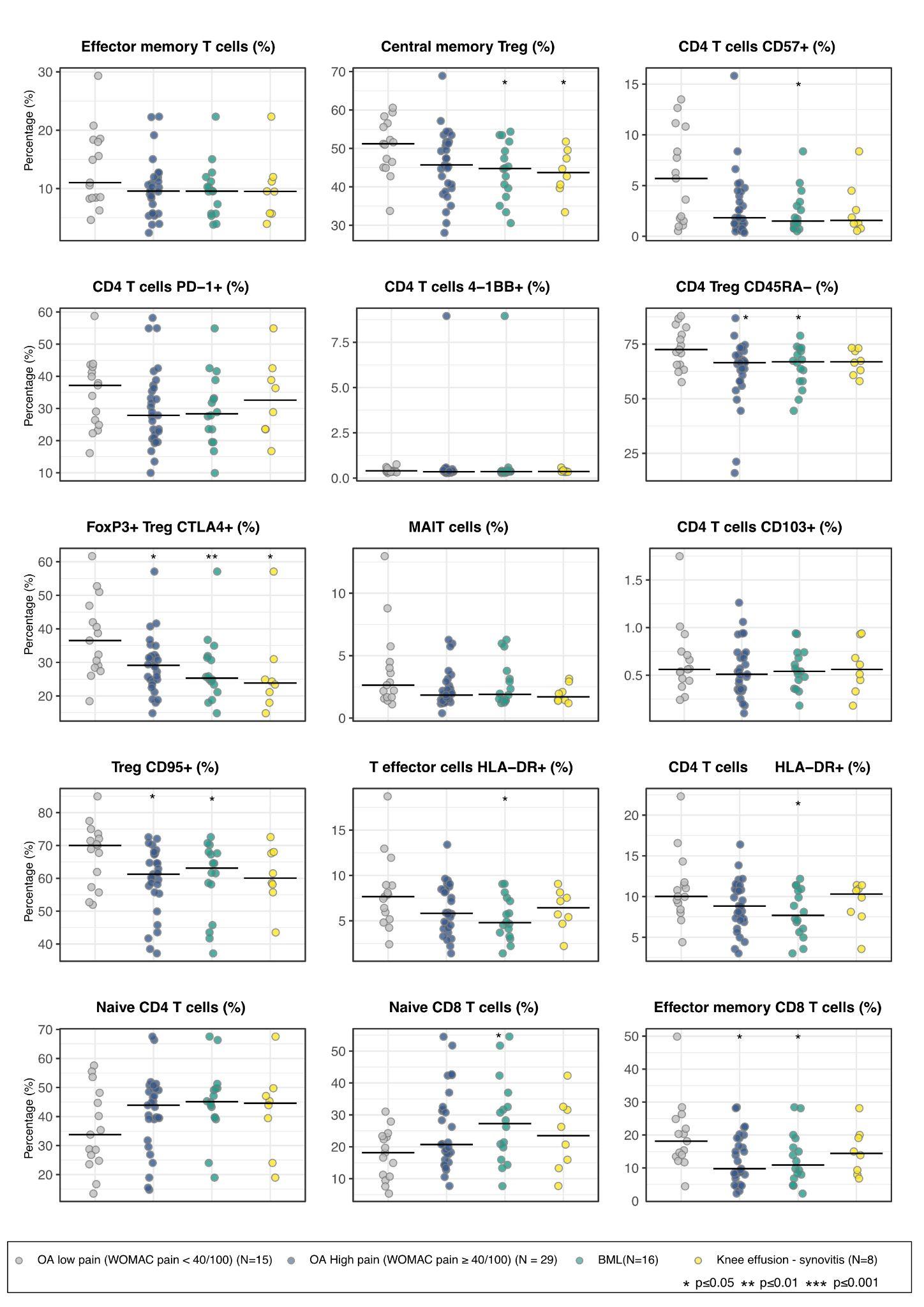

CD, Cluster of differentiation; HLA-DR, Human leukocyte antigen-DR isotype; MAIT, Mucosal-associated invariant T; PD-1, Programmed Cell Death Protein 1; WOMAC, Western Ontario and McMaster universities arthritis index.

**Table S1 | Inclusion and Exclusion criteria for OA patients in the Transimmunom study.**

| **Inclusion criteria** | - Uni or bilateral radiographic knee OA with a KL score of 2 or 3 (ACR criteria) - Patients ≥ 35 years old |
| --- | --- |
| **Exclusion criteria** | - KL score ≥ 4 - Inflammatory rheumatic disease (rheumatoid arthritis, spondyloarthritis, psoriasis) - Crystal induced arthropathies (chondrocalcinosis, gout) - Secondary OA (genetic disorder, hemochromatosis, avascular necrosis) - Contre-indication to X-rays - Pregnancy |

KL, Kellgren Lawrence; OA, Osteoarthritis

**Table S2 | Cytokines and Panel measured in the Transimmunom study.** Cytokines were measured using four Luminex panels including Cytokines-Chemokines kit (CC-CK), Highly sensitive kit (HS), Receptor kit, and TH17 kit. Measurements were expressed in MFI (mean fluorescence intensity) and included 62 cytokines.

| **Panel** | **Cytokines abbreviation** | **Cytokines** |
| --- | --- | --- |
| CCCK | EGF | Epidermal Growth Factor |
| CC-CK | Eotaxin | Eotaxin |
| CC-CK | FGF2 | Fibroblast Growth Factor 2 |
| CC-CK | Fractalkine | Fractalkine |
| CC-CK | GCSF | Granulocyte Colony-Stimulating Factor |
| CC-CK | GMCSF | Granulocyte-Macrophage Colony-Stimulating Factor |
| CC-CK | GRO | Growth-Regulated Oncogene |
| CC-CK | IFNa2 | Interferon Alpha 2 |
| CC-CK | IFNg | Interferon Gamma |
| CC-CK | IL10 | Interleukin 10 |
| CC-CK | IL12p40 | Interleukin 12p40 |
| CC-CK | IL12p70 | Interleukin 12p70 |
| CC-CK | IL13 | Interleukin 13 |
| CC-CK | IL15 | Interleukin 15 |
| CC-CK | IL1a | Interleukin 1 Alpha |
| CC-CK | IL1b | Interleukin 1 Beta |
| CC-CK | IL1RA | Interleukin 1 Receptor Antagonist |
| CC-CK | IL2 | Interleukin 2 |
| CC-CK | IL3 | Interleukin 3 |
| CC-CK | IL4 | Interleukin 4 |
| CC-CK | IL5 | Interleukin 5 |
| CC-CK | IL6 | Interleukin 6 |
| CC-CK | IL7 | Interleukin 7 |
| CC-CK | IL8 | Interleukin 8 |
| CC-CK | IL9 | Interleukin 9 |
| CC-CK | IP10 | Interferon-gamma-Inducible Protein 10 |
| CC-CK | MCP1 | Monocyte Chemoattractant Protein 1 |
| CC-CK | MCP3 | Monocyte Chemoattractant Protein 3 |
| CC-CK | MDC | Macrophage-Derived Chemokine |
| CC-CK | MIP1a | Macrophage Inflammatory Protein 1 Alpha |
| CC-CK | MIP1b | Macrophage Inflammatory Protein 1 Beta |
| CC-CK | sCD40L | Soluble CD40L |
| CC-CK | TGFa | Transforming Growth Factor Alpha |
| CC-CK | TNFa | Tumor Necrosis Factor Alpha |
| CC-CK | TNFb | Tumor Necrosis Factor Beta |
| CC-CK | VEGF | Vascular Endothelial Growth Factor |
| Receptor | sCD30 | Soluble CD30 |
| Receptor | sEGFR | Epidermal Growth Factor Receptor |
| Receptor | sgp130 | Soluble gp130 |
| Receptor | sIL1R1 | Interleukin 1 Receptor Type 1 |
| Receptor | sIL1R2 | Interleukin 1 Receptor Type 2 |
| Receptor | sIL2Ra | Interleukin 2 Receptor Alpha |
| Receptor | sIL4R | Interleukin 4 Receptor |
| Receptor | sIL6R | Interleukin 6 Receptor |
| Receptor | sRAGE | Receptor for Advanced Glycation End Products |
| Receptor | sTNFR1 | Tumor Necrosis Factor Receptor Type 1 |
| Receptor | sTNFR2 | Tumor Necrosis Factor Receptor Type 2 |
| Receptor | sVEGFR1 | Vascular Endothelial Growth Factor Receptor 1 |
| Receptor | sVEGFR2 | Vascular Endothelial Growth Factor Receptor 2 |
| Receptor | sVEGFR3 | Vascular Endothelial Growth Factor Receptor 3 |
| TGFBeta | TGFB1 | Transforming Growth Factor Beta 1 |
| TH17 | IL17A | Interleukin 17A |
| TH17 | IL17E | Interleukin 17E |
| TH17 | IL17F | Interleukin 17F |
| TH17 | IL21 | Interleukin 21 |
| TH17 | IL22 | Interleukin 22 |
| TH17 | IL23 | Interleukin 23 |
| TH17 | IL27 | Interleukin 27 |
| TH17 | IL28A | Interleukin 28A |
| TH17 | IL31 | Interleukin 31 |
| TH17 | IL33 | Interleukin 33 |
| TH17 | MIP3a | Macrophage Inflammatory Protein 3 alpha |

CC-CK, Cytokines - Chemokines kit; Chemokine (C-X3-C motif) ligand 1 (CX3CL1); EGF, Epidermal Growth Factor; Eotaxin, Eosinophil Chemotactic Protein; FGF2, Fibroblast Growth Factor 2; Fractalkine, GCSF, Granulocyte Colony-Stimulating Factor; GMCSF, Granulocyte-Macrophage Colony-Stimulating Factor; GRO, Growth-Regulated Oncogene; IFN, Interferon; IL, Interleukin; IL-1RA, Interleukin 1 Receptor Antagonist; IP-10, Interferon Gamma-Induced Protein 10; MCP, Monocyte Chemoattractant Protein 1; MDC, Macrophage-Derived Chemokine; MIP, Macrophage Inflammatory Protein, sCD40L, Soluble CD40 Ligand; sEGFR, Soluble Epidermal Growth Factor Receptor; sgp130, Soluble Glycoprotein 130; sIL-1R, Soluble Interleukin 1 Receptor; sIL-2Ra, Soluble Interleukin 2 Receptor Alpha; sIL-4R, Soluble Interleukin 4 Receptor; sIL-6R, Soluble Interleukin 6 Receptor; sRAGE, Soluble Receptor for Advanced Glycation End-products; sTNFR, Soluble Tumor Necrosis Factor Receptor; sVEGFR, Soluble Vascular Endothelial Growth Factor Receptor; TGF-alpha, Transforming Growth Factor Alpha; TGF-beta1, Transforming Growth Factor Beta 1; TH, T helper; TNF, Tumor Necrosis Factor; VEGF, Vascular Endothelial Growth Factor.

**Table S3 | Spearman correlation of cytokines significantly associated with WOMAC pain and DN4.** Significant correlations between cell subsets and symptoms were represented using a threshold of p ≤ 0.05 and |r| ≥ 0.2.

| **Symptoms** | **Cytokines** | **Panel** | **r** | **p-value** |
| --- | --- | --- | --- | --- |
| WOMAC pain | IL22 | TH17 | -0.33 | 0.028 |
| WOMAC pain | sIL2Ra | Receptor | 0.31 | 0.021 |
| WOMAC pain | sTNFR1 | Receptor | 0.33 | 0.027 |
| WOMAC pain | sTNFR2 | Receptor | 0.33 | 0.031 |
| DN4 | IL27 | TH17 | -0.35 | 0.021 |
| DN4 | IL4 | CC-CK | -0.33 | 0.028 |

CC-CK, chemokines, cytokines kit; DN4, Douleur neuropathique 4; IL, Interleukin; sTNFR, soluble Tumor necrosis factor receptor; WOMAC, Western Ontario and McMaster universities arthritis index

**Table S4 | Deep immunophenotyping cells populations and panels.** Cell populations were measured across 13 panels, including a general panel, B cell panel, NK-NCR cell panel, T cell activation and T cell naive/memory panel, MAIT-NKT cell panel, T cell migration panel, CD4 T cell polarization panel, T cell activation panel, Treg function panel, DC/monocyte panel, Treg phenotype panel, and Myeloid-ILCs panel. The measurements were expressed in percentage (%) and include 269 cell populations across these panels.

| **Panel** | **Panel name** | **Variable name** | **Cell type** |
| --- | --- | --- | --- |
| Panel 01 | B cells | B cells (%) | B cells |
| Panel 01 | DC/monocytes | Monocytes (%) | Monocytes |
| Panel 01 | DC/monocytes | Classical Monocytes (%) | Monocytes |
| Panel 01 | DC/monocytes | Transient Monocytes (%) | Monocytes |
| Panel 01 | DC/monocytes | Resident Monocytes (%) | Monocytes |
| Panel 01 | Lymphoid | Lymphocytes (%) | Lymphocytes |
| Panel 01 | Memory T cells | Naive CD4 T cells (%) | CD4 T cells |
| Panel 01 | Myeloid - ILCs | Neutrophils/Eosinophils (%) | Neutrophils |
| Panel 01 | Myeloid - ILCs | Eosinophils (%) | Eosinophils |
| Panel 01 | Myeloid - ILCs | Neutrophils (%) | Neutrophils |
| Panel 01 | NK - NCR cells | NK cells (%) | NK cells |
| Panel 01 | NK - NCR cells | CD56dim CD16+ NK cells (%) | NK cells |
| Panel 01 | NK - NCR cells | CD56bright NK cells (%) | NK cells |
| Panel 01 | NK - NCR cells | CD56bright CD16+ NK cells (%) | NK cells |
| Panel 01 | T cells | T cells (%) | T cells |
| Panel 01 | T cells | NKT-like cells (%) | NKT cells |
| Panel 01 | T cells | CD4 T cells (%) | CD4 T cells |
| Panel 01 | T cells | CD8 T cells (%) | CD8 T cells |
| Panel 01 | T cells naive/memory | Central memory CD4 T cells (%) | CD4 T cells |
| Panel 01 | T cells naive/memory | Effector memory CD4 T cells (%) | CD4 T cells |
| Panel 01 | T cells naive/memory | TEMRA CD4+ cells (%) | CD4 T cells |
| Panel 01 | T cells naive/memory | Naive CD8 T cells (%) | CD8 T cells |
| Panel 01 | T cells naive/memory | Central memory CD8 T cells (%) | CD8 T cells |
| Panel 01 | T cells naive/memory | Effector memory CD8 T cells (%) | CD8 T cells |
| Panel 01 | T cells naive/memory | TEMRA CD8 T cells (%) | CD8 T cells |
| Panel 01 | Treg phenotypes | Treg CD25+ CD127+ cells (%) | CD4 T cells |
| Panel 02 | B cells | Naïve B cells (%) | B cells |
| Panel 02 | B cells | Unswitched memory B cells (%) | B cells |
| Panel 02 | B cells | Switched memory B cells (%) | B cells |
| Panel 02 | B cells | IgM+ IgD switched B cells (%) | B cells |
| Panel 02 | B cells | Plasmablast (%) | B cells |
| Panel 02 | B cells | Transitional B cells (%) | B cells |
| Panel 02 | B cells | CD21low B cells (%) | B cells |
| Panel 02 | B cells | CD10+ B cells (%) | B cells |
| Panel 02 | B cells | CD5+ B cells (%) | B cells |
| Panel 02 | B cells | Activated B cells CD32- (%) | B cells |
| Panel 03 | NK - NCR cells | CD56dim NKp44+ cells (%) | NK cells |
| Panel 03 | NK - NCR cells | CD56bright NKp44+ cells (%) | NK cells |
| Panel 03 | NK - NCR cells | CD56dim NKp30+ cells (%) | NK cells |
| Panel 03 | NK - NCR cells | CD56bright NKp30+ cells (%) | NK cells |
| Panel 03 | NK - NCR cells | CD56dim NKp46+ cells (%) | NK cells |
| Panel 03 | NK - NCR cells | CD56brightNKp46+ cells (%) | NK cells |
| Panel 03 | NK - NCR cells | CD56dim NKG2D+ cells (%) | NK cells |
| Panel 03 | NK - NCR cells | CD56bright NKG2D+ cells (%) | NK cells |
| Panel 03 | NK - NCR cells | CD56dim CD8+ NK cells (%) | NK cells |
| Panel 03 | NK - NCR cells | CD56bright CD8+ NK cells (%) | NK cells |
| Panel 03 | NK - NCR cells | CD56dim HLA-DR+ NK cells (%) | NK cells |
| Panel 03 | NK - NCR cells | CD56bright HLA-DR+ NK cells (%) | NK cells |
| Panel 03 | NK - NCR cells | NKG2D+ CD8 T cells (%) | CD8 T cells |
| Panel 03 | NK - NCR cells | HLA-DR+ CD8 T cells (%) | CD8 T cells |
| Panel 03 | NK - NCR cells | TCRgd T cells (%) | CD8 T cells |
| Panel 03 | NK - NCR cells | TCRgd CD16+ T cells (%) | TCRgd T cells |
| Panel 03 | NK - NCR cells | TCRgd NKG2D+ T cells (%) | TCRgd T cells |
| Panel 03 | NK - NCR cells | TCRgd HLA-DR+ T cells (%) | TCRgd T cells |
| Panel 03 | NK - NCR cells | TCRgd CD8+ T cells (%) | TCRgd T cells |
| Panel 03 | NK - NCR cells | NKT-like CD8+ (%) | NKT cells |
| Panel 03 | NK - NCR cells | NKT-like CD16+ cells (%) | NKT cells |
| Panel 03 | NK - NCR cells | NKT-like HLA-DR+ cells (%) | NKT cells |
| Panel 03 | NK - NCR cells | NKT-like NKG2D+ cells (%) | NKT cells |
| Panel 04 | T cells activation | CD8 T cells CD95+ (%) | CD8 T cells |
| Panel 04 | T cells activation | T effector cells CD95+ (%) | CD4 T cells |
| Panel 04 | T cells activation | Treg CD95+ (%) | CD4 T cells |
| Panel 04 | T cells activation | CD4 T cells CD95+ (%) | CD4 T cells |
| Panel 04 | T cells activation | CD8 T cells HLA-DR+ (%) | CD8 T cells |
| Panel 04 | T cells activation | T effector cells HLA-DR+ (%) | CD4 T cells |
| Panel 04 | T cells activation | Treg HLA-DR+ (%) | CD4 T cells |
| Panel 04 | T cells activation | CD4 T cells HLA-DR+ (%) | CD4 T cells |
| Panel 04 | T cells activation | CD8 ICOS+ (%) | CD8 T cells |
| Panel 04 | T cells activation | T effector cells ICOS+ (%) | CD4 T cells |
| Panel 04 | T cells activation | Treg ICOS+ (%) | CD4 T cells |
| Panel 04 | T cells activation | CD4 T cells ICOS+ (%) | CD4 T cells |
| Panel 04 | T cells naive/memory | CD4 T effector (%) | CD4 T cells |
| Panel 04 | T cells naive/memory | Naive T effector cells (%) | CD4 T cells |
| Panel 04 | T cells naive/memory | Central effector memory T cells (%) | CD4 T cells |
| Panel 04 | T cells naive/memory | Effector memory T cells (%) | CD4 T cells |
| Panel 04 | T cells naive/memory | TEMRA cells (%) | CD4 T cells |
| Panel 04 | T cells naive/memory | Naive T reg (%) | CD4 T cells |
| Panel 04 | T cells naive/memory | Central memory Treg (%) | CD4 T cells |
| Panel 04 | T cells naive/memory | Effector memory Treg (%) | CD4 T cells |
| Panel 04 | T cells naive/memory | TEMRA Treg (%) | CD4 T cells |
| Panel 05 | MAIT - NKT cells | NKT-like cells (%) | NKT cells |
| Panel 05 | MAIT - NKT cells | NKT TCR-Va7.2+ cells (%) | NKT cells |
| Panel 05 | MAIT - NKT cells | MAIT cells (%) | MCD4 T cells |
| Panel 07 | T cells migration | CD4 T cells CLA+ (%) | CD8 T cells |
| Panel 07 | T cells migration | CD8 T cells CLA+ (%) | CD4 T cells |
| Panel 07 | T cells migration | CD4 T cells CD49a+ (%) | CD8 T cells |
| Panel 07 | T cells migration | CD8 T cells CD49a+ (%) | CD4 T cells |
| Panel 07 | T cells migration | CD4 T cells CD103+ (%) | CD8 T cells |
| Panel 07 | T cells migration | CD8 T cells CD103+ (%) | CD4 T cells |
| Panel 07 | T cells migration | CD4 T cells a4+B7+ (%) | CD8 T cells |
| Panel 07 | T cells migration | CD8 T cells a4+B7+ (%) | CD4 T cells |
| Panel 07 | T cells migration | CD4 T cells a4+B7- (%) | CD8 T cells |
| Panel 07 | T cells migration | CD8 T cells a4+B7- (%) | CD8 T cells |
| Panel 08 | CD4 T cells polarization | TEMRA Th2 cells (%) | CD4 T cells |
| Panel 08 | CD4 T cells polarization | TEMRA Th17 cells (%) | CD4 T cells |
| Panel 08 | CD4 T cells polarization | TEMRA Th22 cells (%) | CD4 T cells |
| Panel 08 | CD4 T cells polarization | T helper Th2 cells (%) | CD4 T cells |
| Panel 08 | CD4 T cells polarization | T helper Th17 cells (%) | CD4 T cells |
| Panel 08 | CD4 T cells polarization | T helper Th1 cells (%) | CD4 T cells |
| Panel 08 | CD4 T cells polarization | T helper Th9-Th17.1 cells (%) | CD4 T cells |
| Panel 08 | CD4 T cells polarization | T helper Th22 cells (%) | CD4 T cells |
| Panel 08 | CD4 T cells polarization | Th1 memory cells (%) | CD4 T cells |
| Panel 08 | CD4 T cells polarization | Th2 memory cells (%) | CD4 T cells |
| Panel 08 | CD4 T cells polarization | Th17 memory cells (%) | CD4 T cells |
| Panel 08 | CD4 T cells polarization | Th22 memory cells (%) | CD4 T cells |
| Panel 08 | CD4 T cells polarization | TEMRA Th1 cells (%) | CD4 T cells |
| Panel 09 | T cells activation | CD4 T cells CD57+ (%) | CD4 T cells |
| Panel 09 | T cells activation | CD8 T cells CD57+ (%) | CD8 T cells |
| Panel 09 | T cells activation | CD4 T cells CD69+ (%) | CD4 T cells |
| Panel 09 | T cells activation | CD8 T cells CD69+ (%) | CD8 T cells |
| Panel 09 | T cells activation | CD4 T cells PD-1+ (%) | CD4 T cells |
| Panel 09 | T cells activation | CD8 T cells PD-1+ (%) | CD8 T cells |
| Panel 09 | T cells activation | CD4 T cells 4-1BB+ (%) | CD4 T cells |
| Panel 09 | T cells activation | CD8 T cells 4-1BB+ (%) | CD8 T cells |
| Panel 09 | T cells activation | CD4 T cells OX40+ (%) | CD4 T cells |
| Panel 09 | T cells activation | CD8 T cells OX40+ (%) | CD8 T cells |
| Panel 09 | T cells activation | CD4 T cells CD5hi (%) | CD4 T cells |
| Panel 09 | T cells activation | CD8 T cells CD5hi (%) | CD8 T cells |
| Panel 10 | Treg function | CD4 Treg CD39+ (%) | CD4 T cells |
| Panel 10 | Treg function | CD4 Treg GITR+ (%) | CD4 T cells |
| Panel 10 | Treg function | CD4 Treg LAP+ (%) | CD4 T cells |
| Panel 10 | Treg function | CD4 Treg LAG3+ (%) | CD4 T cells |
| Panel 10 | Treg function | CD4 Treg CD45RA- (%) | CD4 T cells |
| Panel 11 | DC/monocytes | pDCs (%) | DC/monocytes |
| Panel 11 | DC/monocytes | mDC1 (%) | DC/monocytes |
| Panel 11 | DC/monocytes | mDC2 (%) | DC/monocytes |
| Panel 11 | DC/monocytes | mDC CCR5+ (%) | DC/monocytes |
| Panel 11 | DC/monocytes | mDC (%) | DC/monocytes |
| Panel 11 | DC/monocytes | Basophils (%) | DC/monocytes |
| Panel 12 | Treg phenotypes | FoxP3+ Treg (%) | CD4 T cells |
| Panel 12 | Treg phenotypes | FoxP3+ Treg CXCR5+ (%) | CD4 T cells |
| Panel 12 | Treg phenotypes | FoxP3+ Treg CXCR5+ (%) | CD4 T cells |
| Panel 12 | Treg phenotypes | FoxP3+ Treg CTLA4+ (%) | CD4 T cells |
| Panel 13 | Myeloid - ILCs | Innate lymphoid cells (%) | ILCs |
| Panel 13 | Myeloid - ILCs | Innate lymphoid cells ILC2 (%) | ILCs |
| Panel 13 | Myeloid - ILCs | Innate lymphoid cells ILC1 (%) | ILCs |
| Panel 13 | Myeloid - ILCs | Innate lymphoid cells ILC3 (%) | ILCs |

CCR Chemokine receptor; CD, Cluster of differentiation; CLA, Cutaneous leukocyte-associated antigen; CTLA-4, Cytotoxic T-lymphocyte–associated antigen; DC dendritics cells, FoxP3, Forkhead box P3; GITR, Glucocorticoid-Induced TNFR-Related; HLA-DR, Human leukocyte antigen-DR isotype; ICOS, Inducible costimulator; ILC, Innate lymphoid cells; Ig, Immunoglobulin; LAG-3, Lymphocyte activation gene 3; LAP, LC3-associated phagocytosis, MAIT, Mucosal-associated invariant T; NK, Natural killer; NKG2D, Natural killer group 2 member D; PD-1, Programmed Cell Death Protein; TCR, T-cell receptor, TEMRA, Terminally differentiated effector memory cell; Treg, T regulatory cell

**Table S5 | Spearman correlation of cell population significantly associated with WOMAC pain and DN4.** Significant correlations between cell subsets and symptoms were represented using a threshold of p ≤ 0.05 and |r| ≥ 0.2.

| **Symptoms** | **Cell population** | **Panel** | **r** | **p-value** |  |
| --- | --- | --- | --- | --- | --- |
| WOMAC pain | Effector memory T cells (%) | T cells naive/memory | -0.36 | 0.018 | * |
| WOMAC pain | Central memory Treg (%) | T cells naive/memory | -0.33 | 0.027 | * |
| WOMAC pain | CD4 T cells CD57+ (%) | T cells activation | -0.34 | 0.023 | * |
| WOMAC pain | CD4 T cells PD-1+ (%) | T cells activation | -0.37 | 0.012 | * |
| WOMAC pain | CD4 T cells 4-1BB+ (%) | T cells activation | -0.33 | 0.029 | * |
| WOMAC pain | CD4 Treg CD45RA- (%) | Treg function | -0.39 | 0.009 | ** |
| WOMAC pain | FoxP3+ Treg CTLA4+ (%) | Treg phenotypes | -0.41 | 0.006 | ** |
| WOMAC pain | MAIT cells (%) | MAIT - NKT cells | -0.31 | 0.039 | * |
| WOMAC pain | CD4 T cells CD103+ (%) | T cells migration | -0.3 | 0.046 | * |
| WOMAC pain | Treg CD95+ (%) | T cells activation | -0.33 | 0.028 | * |
| WOMAC pain | T effector cells HLA-DR+ (%) | T cells activation | -0.36 | 0.017 | * |
| WOMAC pain | CD4 T cells HLA-DR+ (%) | T cells activation | -0.36 | 0.018 | * |
| WOMAC pain | Naive CD4 T cells (%) | Memory T cells | 0.36 | 0.017 | * |
| WOMAC pain | Naive CD8 T cells (%) | T cells naive/memory | 0.37 | 0.013 | * |
| DN4 | Neutrophils/Eosinophils (%) | Myeloid - ILCs | -0.33 | 0.035 | * |
| DN4 | Lymphocytes (%) | Lymphoid | 0.45 | 0.003 | ** |
| DN4 | CD56dim NKp44+ cells (%) | NK - NCR cells | 0.48 | 0.001 | ** |
| DN4 | CD56bright NKp30+ cells (%) | NK - NCR cells | 0.38 | 0.013 | * |
| DN4 | CD56bright CD8+ NK cells (%) | NK - NCR cells | 0.35 | 0.023 | * |
| DN4 | Effector memory Treg (%) | T cells naive/memory | -0.36 | 0.018 | * |
| DN4 | FoxP3+ Treg CXCR5+ (%) | Treg phenotypes | 0.39 | 0.011 | * |
| DN4 | FoxP3+ Treg CTLA4+ (%) | Treg phenotypes | -0.36 | 0.02 | * |
| DN4 | Treg CD95+ (%) | T cells activation | -0.35 | 0.023 | * |
| DN4 | Eosinophils (%) | Myeloid - ILCs | 0.37 | 0.017 | * |
| DN4 | Neutrophils (%) | Myeloid - ILCs | -0.38 | 0.012 | * |
| DN4 | Effector memory CD4 T cells (%) | T cells naive/memory | -0.32 | 0.041 | * |

CD, Cluster differentiation; DN4, Douleur neuropathique 4; HLA-DR, Human leukocyte antigen-DR isotype; MAIT, Mucosal-associated invariant T cells; NK, Natural Killer; PD-1, Programmed Cell Death Protein 1; TCR, T cell receptor; TEMRA,T effector memory re-expressing CD45RA; WOMAC, Western Ontario and McMaster universities arthritis index

**Table S6 | Differentially expressed genes in Tregs of patients with low (N=6) and high pain intensity in Treg (N=28).** (|log2 FC| ≥ log2(1.6), p-value ≤ 0.01, baseMean expression ≥ 8).

|  | **Base mean** | **log2(FC)** | **p-value** |
| --- | --- | --- | --- |
| CAVIN3 | 11.65 | -1.58 | 0.0092 |
| ADAM12 | 143.37 | -1.34 | 0.0003 |
| MRC2 | 33.04 | -1.28 | 0.0061 |
| GPRIN1 | 26.63 | -1.25 | 0.0004 |
| GDF7 | 11.32 | -1.23 | 0.0062 |
| IL23R | 32.8 | -1.17 | 0.0042 |
| CACNB3 | 25.86 | -1.15 | 0.003 |
| COPG2 | 48.75 | -1.05 | 0.0009 |
| HYAL2 | 26.42 | -1 | 0.0076 |
| NPR2 | 26.03 | -0.99 | 0.0073 |
| CELSR3 | 62.63 | -0.95 | 0.0086 |
| TNFSF13B | 140.35 | -0.9 | 0.0075 |
| RORC | 152.87 | -0.84 | 0.0002 |
| DHTKD1 | 295.15 | -0.8 | 0.0002 |
| SKAP2 | 394.78 | -0.8 | 0.0022 |
| ZNF213 | 48.49 | -0.8 | 0.0053 |
| CEP55 | 106.83 | -0.77 | 0.0088 |
| ZSCAN20 | 22.72 | -0.77 | 0.0044 |
| TRAIP | 53.87 | -0.76 | 0.0012 |
| CYFIP1 | 80.06 | -0.73 | 0.006 |
| EEF2KMT | 95.11 | -0.71 | 0.0082 |
| MEOX1 | 180.16 | -0.71 | 0.0001 |
| MAN2C1 | 1195.76 | -0.69 | 0.0012 |
| ZNF268 | 388.69 | 0.69 | 0.002 |
| ZNF546 | 112.99 | 0.71 | 0.0003 |
| RYBP | 245.51 | 0.72 | 0.0011 |
| FCGRT | 393.95 | 0.79 | 0.0001 |
| RGS12 | 277.16 | 0.79 | 0.001 |
| SLC19A3 | 34.6 | 0.79 | 0.0074 |
| USP18 | 124.53 | 0.82 | 0.0035 |
| ANKRD42 | 93.6 | 0.87 | 1.01E-05 |
| SEC14L2 | 192.61 | 0.87 | 0.0044 |
| BOLA2B | 96.98 | 0.97 | 0.0052 |
| CYP2D6 | 26.53 | 1.01 | 0.0063 |
| F2RL1 | 35.09 | 1.15 | 0.0061 |
| RGPD6 | 283.56 | 1.2 | 0.0085 |
| GSTM3 | 261.66 | 1.21 | 0.0067 |
| IFITM3 | 232.48 | 1.25 | 0.007 |
| IFI44L | 82.85 | 1.31 | 0.0015 |
| RAB31 | 48.27 | 1.36 | 0.0093 |
| CD244 | 27.9 | 1.37 | 0.0076 |
| PPFIBP2 | 48.66 | 1.39 | 0.0044 |
| WWC1 | 21.54 | 1.41 | 0.0087 |
| GPR150 | 16.29 | 1.42 | 0.0004 |
| IL31RA | 15.84 | 1.42 | 0.0078 |
| SIGLEC14 | 23.66 | 1.45 | 0.0094 |
| CD9 | 66.9 | 1.54 | 0.0005 |
| CFD | 62.32 | 1.65 | 0.0067 |
| LILRA2 | 41.67 | 1.71 | 0.0039 |
| TSKS | 16.59 | 1.83 | 0.0034 |
| KCTD15 | 11.43 | 1.87 | 0.0084 |
| NEBL | 69 | 1.88 | 0.0022 |
| PYGL | 14.35 | 1.91 | 0.0033 |
| MPIG6B | 27.92 | 2.04 | 0.0023 |
| GATA2 | 26.84 | 2.05 | 0.0053 |
| MAMLD1 | 10.92 | 2.21 | 0.0014 |
| CLU | 113.45 | 2.37 | 0.0045 |
| EHD2 | 10.24 | 2.42 | 0.0006 |
| NLRP3 | 35.24 | 2.59 | 0.0038 |
| IL1RL1 | 16.94 | 2.62 | 0.0007 |
| CACHD1 | 9.67 | 2.7 | 0.0015 |
| CLC | 26.56 | 2.85 | 0.0028 |
| FCER1A | 111.55 | 2.99 | 0.0004 |
| FPR1 | 36.94 | 3.21 | 0.0048 |
| TCN1 | 13.34 | 3.25 | 0.0002 |
| BCL2L2-PABPN1 | 9.09 | 4.11 | 0.0015 |
| PTGS1 | 21.49 | 5.32 | 8.41E-06 |
| H2AC19 | 31.72 | 7.79 | 1.01E-05 |
| UPK3BL2 | 26.56 | 21.83 | 6.72E-12 |

**Table S7 | Differentially expressed genes in Teff of patients with low (N=14) and high pain intensity in Treg (N=28).** (|log2 FC| ≥ log2(1.6), p-value ≤ 0.01 baseMean expression ≥ 8).

|  | **Base mean** | **Log2(FC)** | **p-value** |
| --- | --- | --- | --- |
| CD14 | 28.19 | -1.7 | 0.0001 |
| PTGS1 | 81.58 | -1.66 | 0.001 |
| C2orf88 | 29.25 | -1.4 | 0.0058 |
| CD1C | 26.52 | -1.35 | 0.0086 |
| KLRC4-KLRK1 | 31.97 | -1.28 | 0.0068 |
| CA2 | 23.34 | -1.26 | 0.0067 |
| MS4A7 | 239.1 | -1.26 | 0.0033 |
| CDKN1C | 29 | -1.24 | 0.0001 |
| FCN1 | 250.15 | -1.24 | 0.0045 |
| FCGR3A | 1214.68 | -1.17 | 0.0082 |
| PLBD1 | 18.15 | -1.12 | 0.0058 |
| RAMP1 | 22.26 | -1.09 | 0.007 |
| KCNMB3 | 35.46 | -1.07 | 0.0049 |
| CLEC7A | 101.96 | -1.06 | 0.0041 |
| TNFRSF11A | 30.4 | -1.05 | 0.0015 |
| TYROBP | 294.68 | -1.05 | 0.0077 |
| CD86 | 56.02 | -1.01 | 0.0064 |
| CEBPD | 129.8 | -1 | 0.0026 |
| LMO2 | 29.17 | -0.99 | 0.0031 |
| MARCHF1 | 113.22 | -0.98 | 0.0089 |
| METTL7A | 118.51 | -0.98 | 0.0005 |
| TNFSF13B | 170.2 | -0.98 | 0.0013 |
| GPRIN1 | 19.8 | -0.97 | 0.0096 |
| CPVL | 71.57 | -0.96 | 0.0026 |
| CST3 | 249.12 | -0.96 | 0.0054 |
| GUCY1A1 | 29.91 | -0.96 | 0.0046 |
| KYNU | 156.38 | -0.93 | 0.0006 |
| TNFSF13 | 40.07 | -0.92 | 0.0036 |
| IFI30 | 517.51 | -0.88 | 0.0094 |
| CEBPA | 43.53 | -0.86 | 0.0082 |
| SLC46A1 | 43.19 | -0.86 | 0.0051 |
| NEFH | 19.35 | -0.84 | 0.0056 |
| FGL2 | 160.48 | -0.83 | 0.0078 |
| KCNE3 | 14.92 | -0.82 | 0.0049 |
| PCDHGB6 | 15.89 | -0.81 | 0.0093 |
| SECTM1 | 73.33 | -0.81 | 0.007 |
| PRR5L | 157.54 | -0.78 | 0.0001 |
| CDK2AP1 | 91.52 | -0.77 | 0.001 |
| ALOX12 | 34.57 | -0.75 | 0.0007 |
| NFIL3 | 109.49 | -0.75 | 0.0018 |
| DAPK1 | 56.96 | -0.74 | 0.0074 |
| PDLIM1 | 120.64 | -0.74 | 0.0073 |
| PLEK | 754.09 | -0.72 | 0.0035 |
| SIGLEC14 | 48.89 | -0.71 | 0.0096 |
| SKAP2 | 286.12 | -0.71 | 0.0054 |
| RHOC | 313.35 | -0.7 | 0.0022 |
| MBD2 | 235.53 | -0.69 | 0.0013 |
| CMC1 | 232.99 | -0.68 | 0.0002 |
| FUT7 | 90.78 | -0.68 | 0.0075 |
| ARHGEF5 | 65.67 | 0.71 | 0.0027 |
| FLT4 | 107 | 0.71 | 0.0094 |
| POLR2J2 | 770.56 | 0.73 | 0.0042 |
| ADAMTSL2 | 33.33 | 0.75 | 0.0009 |
| FCGBP | 116.19 | 0.75 | 0.0069 |
| CACNA1H | 73.72 | 0.84 | 0.0061 |
| WDR97 | 70.24 | 1.07 | 0.0016 |
| GCOM1 | 37.22 | 1.11 | 0.0011 |
| NPIPA7 | 28.47 | 1.3 | 0.0042 |
| ALAS2 | 227.96 | 1.62 | 0.0066 |
| EPB42 | 18.77 | 1.95 | 0.0012 |

**Table S8 | Non-parametric Wilcoxon analysis comparing cell populations in patients with WOMAC pain score ≥40/100, BML (N=16), knee effusion (N=8) to patients with low pain intensity.** A significant p-value was defined with a threshold of ≤ 0.05.

| **Cell population** | **Group 1** | **Group 2** | **p** |  |
| --- | --- | --- | --- | --- |
| Effector memory T cells (%) | OA low pain  (WOMAC pain < 40/100) (N=15) | OA high pain (WOMAC pain ≥ 40/100) (N = 29) | 0.173 |  |
| Effector memory T cells (%) | OA ow pain  (WOMAC pain < 40/100) (N=15) | BML (N=16) | 0.138 |  |
| Effector memory T cells (%) | OA low pain  (WOMAC pain < 40/100) (N=15) | Knee effusion (N=8) | 0.325 |  |
| Central memory Treg (%) | OA low pain (WOMAC pain < 40/100) (N=15) | OA high pain (WOMAC pain ≥ 40/100) (N = 29) | 0.066 | . |
| Central memory Treg (%) | OA low pain  WOMAC pain < 40/100) (N=15) | BML (N=16) | 0.027 | * |
| Central memory Treg (%) | OA low pain  (WOMAC pain < 40/100) (N=15) | Knee effusion (N=8) | 0.028 | * |
| CD4 T cells CD57+ (%) | OA low pain  (WOMAC pain < 40/100) (N=15) | OA high pain (WOMAC pain ≥ 40/100) (N = 29) | 0.051 | . |
| CD4 T cells CD57+ (%) | OA low pain  (WOMAC pain < 40/100) (N=15) | BML (N=16) | 0.034 | * |
| CD4 T cells CD57+ (%) | OA low pain  (WOMAC pain < 40/100) (N=15) | Knee effusion (N=8) | 0.149 |  |
| CD4 T cells PD-1+ (%) | OA low pain  (WOMAC pain < 40/100) (N=15) | OA high pain (WOMAC pain ≥ 40/100) (N = 29) | 0.114 |  |
| CD4 T cells PD-1+ (%) | OA low pain  (WOMAC pain < 40/100) (N=15) | BML (N=16) | 0.202 |  |
| CD4 T cells PD-1+ (%) | OA low pain  (WOMAC pain < 40/100) (N=15) | Knee effusion (N=8) | 0.681 |  |
| CD4 T cells 4-1BB+ (%) | OA low pain  (WOMAC pain < 40/100) (N=15) | OA high pain (WOMAC pain ≥ 40/100) (N = 29) | 0.068 | . |
| CD4 T cells 4-1BB+ (%) | OA low pain  (WOMAC pain < 40/100) (N=15) | BML (N=16) | 0.133 |  |
| CD4 T cells 4-1BB+ (%) | OA low pain  (WOMAC pain < 40/100) (N=15) | Knee effusion (N=8) | 0.477 |  |
| CD4 Treg CD45RA- (%) | OA low pain  (WOMAC pain < 40/100) (N=15) | OA high pain (WOMAC pain ≥ 40/100) (N = 29) | 0.021 | * |
| CD4 Treg CD45RA- (%) | OA low pain  (WOMAC pain < 40/100) (N=15) | BML (N=16) | 0.04 | * |
| CD4 Treg CD45RA- (%) | OA low pain  (WOMAC pain < 40/100) (N=15) | Knee effusion (N=8) | 0.149 |  |
| FoxP3+ Treg CTLA4+ (%) | OA low pain  (WOMAC pain < 40/100) (N=15) | OA high pain (WOMAC pain ≥ 40/100) (N = 29) | 0.019 | * |
| FoxP3+ Treg CTLA4+ (%) | OA low pain  (WOMAC pain < 40/100) (N=15) | BML (N=16) | 0.009 | ** |
| FoxP3+ Treg CTLA4+ (%) | OA low pain  (WOMAC pain < 40/100) (N=15) | Knee effusion (N=8) | 0.019 | * |
| MAIT cells (%) | OA low pain  (WOMAC pain < 40/100) (N=15) | OA high pain (WOMAC pain ≥ 40/100) (N = 29) | 0.071 | . |
| MAIT cells (%) | OA low pain  (WOMAC pain < 40/100) (N=15) | BML (N=16) | 0.313 |  |
| MAIT cells (%) | OA low pain  (WOMAC pain < 40/100) (N=15) | Knee effusion (N=8) | 0.1 | . |
| CD4 T cells CD103+ (%) | OA low pain  (WOMAC pain < 40/100) (N=15) | OA high pain (WOMAC pain ≥ 40/100) (N = 29) | 0.428 |  |
| CD4 T cells CD103+ (%) | OA low pain  (WOMAC pain < 40/100) (N=15) | BML (N=16) | 0.649 |  |
| CD4 T cells CD103+ (%) | OA low pain  (WOMAC pain < 40/100) (N=15) | Knee effusion (N=8) | 0.796 |  |
| Treg CD95+ (%) | OA low pain  (WOMAC pain < 40/100) (N=15) | OA high pain (WOMAC pain ≥ 40/100) (N = 29) | 0.018 | * |
| Treg CD95+ (%) | OA low pain  (WOMAC pain < 40/100) (N=15) | BML (N=16) | 0.049 | * |
| Treg CD95+ (%) | OA low pain  (WOMAC pain < 40/100) (N=15) | Knee effusion (N=8) | 0.149 |  |
| T effector cells HLA-DR+ (%) | OA low pain  (WOMAC pain < 40/100) (N=15) | OA high pain (WOMAC pain ≥ 40/100) (N = 29) | 0.175 |  |
| T effector cells HLA-DR+ (%) | OA low pain  (WOMAC pain < 40/100) (N=15) | BML (N=16) | 0.033 | * |
| T effector cells HLA-DR+ (%) | OA low pain  (WOMAC pain < 40/100) (N=15) | Knee effusion (N=8) | 0.357 |  |
| CD4 T cells HLA-DR+ (%) | OA low pain  (WOMAC pain < 40/100) (N=15) | OA high pain (WOMAC pain ≥ 40/100) (N = 29) | 0.12 |  |
| CD4 T cells HLA-DR+ (%) | OA low pain  (WOMAC pain < 40/100) (N=15) | BML (N=16) | 0.049 | * |
| CD4 T cells HLA-DR+ (%) | OA low pain  (WOMAC pain < 40/100) (N=15) | Knee effusion (N=8) | 0.506 |  |
| Naive CD4 T cells (%) | OA low pain  (WOMAC pain < 40/100) (N=15) | OA high pain (WOMAC pain ≥ 40/100) (N = 29) | 0.192 |  |
| Naive CD4 T cells (%) | OA low pain  (WOMAC pain < 40/100) (N=15) | BML (N=16) | 0.093 | . |
| Naive CD4 T cells (%) | OA low pain  (WOMAC pain < 40/100) (N=15) | Knee effusion (N=8) | 0.428 |  |
| Naive CD8 T cells (%) | OA low pain  (WOMAC pain < 40/100) (N=15) | OA high pain (WOMAC pain ≥ 40/100) (N = 29) | 0.088 | . |
| Naive CD8 T cells (%) | OA low pain  (WOMAC pain < 40/100) (N=15) | BML (N=16) | 0.027 | * |
| Naive CD8 T cells (%) | OA low pain  (WOMAC pain < 40/100) (N=15) | Knee effusion (N=8) | 0.213 |  |
| Effector memory CD8 T cells (%) | OA low pain  (WOMAC pain < 40/100) (N=15) | OA high pain (WOMAC pain ≥ 40/100) (N = 29) | 0.022 | * |
| Effector memory CD8 T cells (%) | OA low pain  (WOMAC pain < 40/100) (N=15) | BML (N=16) | 0.044 | * |
| Effector memory CD8 T cells (%) | OA low pain  (WOMAC pain < 40/100) (N=15) | Knee effusion (N=8) | 0.265 |  |
| Cell popiulation | OA low pain  (WOMAC pain < 40/100) (N=15) | group2 | p |  |
| Effector memory T cells (%) | OA low pain  (WOMAC pain < 40/100) (N=15) | OA high pain (WOMAC pain ≥ 40/100) (N = 29) | 0.173 |  |
| Effector memory T cells (%) | OA low pain  (WOMAC pain < 40/100) (N=15) | BML (N=16) | 0.138 |  |
| Effector memory T cells (%) | OA low pain  (WOMAC pain < 40/100) (N=15) | Knee effusion (N=8) | 0.325 |  |
| Central memory Treg (%) | OA low pain  (WOMAC pain < 40/100) (N=15) | OA high pain (WOMAC pain ≥ 40/100) (N = 29) | 0.066 | . |
| Central memory Treg (%) | OA low pain  (WOMAC pain < 40/100) (N=15) | BML (N=16) | 0.027 | * |
| Central memory Treg (%) | OA low pain  (WOMAC pain < 40/100) (N=15) | Knee effusion (N=8) | 0.028 | * |
| CD4 T cells CD57+ (%) | OA low pain  (WOMAC pain < 40/100) (N=15) | OA high pain (WOMAC pain ≥ 40/100) (N = 29) | 0.051 | . |
| CD4 T cells CD57+ (%) | OA low pain  (WOMAC pain < 40/100) (N=15) | BML (N=16) | 0.034 | * |
| CD4 T cells CD57+ (%) | OA low pain  (WOMAC pain < 40/100) (N=15) | Knee effusion (N=8) | 0.149 |  |
| CD4 T cells PD-1+ (%) | OA low pain  (WOMAC pain < 40/100) (N=15) | OA high pain (WOMAC pain ≥ 40/100) (N = 29) | 0.114 |  |
| CD4 T cells PD-1+ (%) | OA low pain  (WOMAC pain < 40/100) (N=15) | BML (N=16) | 0.202 |  |
| CD4 T cells PD-1+ (%) | OA low pain  (WOMAC pain < 40/100) (N=15) | Knee effusion (N=8) | 0.681 |  |
| CD4 T cells 4-1BB+ (%) | OA low pain  (WOMAC pain < 40/100) (N=15) | OA high pain (WOMAC pain ≥ 40/100) (N = 29) | 0.068 | . |
| CD4 T cells 4-1BB+ (%) | OA low pain  (WOMAC pain < 40/100) (N=15) | BML (N=16) | 0.133 |  |

CD, Cluster differentiation; HLA-DR, Human leukocyte antigen-DR isotype; MAIT, Mucosal-associated invariant T cells; NK, Natural Killer; PD-1, Programmed Cell Death Protein 1; TCR, T cell receptor

**Table S9 | Non-parametric Mann-Whitney analysis comparing cytokines in patients with WOMAC pain score ≥40/100 and different pain phenotypes (neuropathic pain (N=10), BML (N=16), knee effusion (N=8)).** For each pain phenotype variable, comparisons were made with groups without neuropathic pain (N=32), without BML and pain (N=23), and without knee effusion (N=31). A significant p-value was defined with a threshold of ≤ 0.05.

| **Cytokines** | **Group 1** | **Group 2** | **p-value** |  |
| --- | --- | --- | --- | --- |
| sIL2Ra | OA low pain  (WOMAC pain < 40/100) (N=15) | OA high pain (WOMAC pain≥ 40/100) (N = 29) | 0.105 |  |
| sIL2Ra | OA low pain  (WOMAC pain < 40/100) (N=15) | BML (N=16) | 0.286 |  |
| sIL2Ra | OA low pain  (WOMAC pain < 40/100) (N=15) | Knee effusion (N=8) | 0.213 |  |
| sTNFR1 | OA low pain  (WOMAC pain < 40/100) (N=15) | OA high pain (WOMAC pain≥ 40/100) (N = 29) | 0.16 |  |
| sTNFR1 | OA low pain  (WOMAC pain < 40/100) (N=15) | BML (N=16) | 0.066 | . |
| sTNFR1 | OA low pain  (WOMAC pain < 40/100) (N=15) | Knee effusion (N=8) | 0.065 | . |
| sTNFR2 | OA low pain  (WOMAC pain < 40/100) (N=15) | OA high pain (WOMAC pain≥ 40/100) (N = 29) | 0.028 | * |
| sTNFR2 | OA low pain  (WOMAC pain < 40/100) (N=15) | BML (N=16) | 0.019 | * |
| sTNFR2 | OA low pain  (WOMAC pain < 40/100) (N=15) | Knee effusion (N=8) | 0.076 | . |
| IL22 | OA low pain  (WOMAC pain < 40/100) (N=15) | OA high pain (WOMAC pain≥ 40/100) (N = 29) | 0.013 | * |
| IL22 | OA low pain  (WOMAC pain < 40/100) (N=15) | BML (N=16) | 0.046 | * |
| IL22 | OA low pain  (WOMAC pain < 40/100) (N=15) | Knee effusion (N=8) | 0.018 | * |
